# DUSP12 promotes cell cycle progression and protects cells from cell death by regulating ZPR9

**DOI:** 10.1101/2025.01.13.632830

**Authors:** Mai Abdusamad, Xiao Guo, Ivan Ramirez, Erick F. Velasquez, Whitaker Cohn, Ankur A. Gholkar, Julian P. Whitelegge, Jorge Z. Torres

## Abstract

Protein phosphatases are critical for regulating cell signaling, cell cycle, and cell fate decisions, and their dysregulation leads to an array of human diseases like cancer. The dual specificity phosphatases (DUSPs) have emerged as important factors driving tumorigenesis and cancer therapy resistance. DUSP12 is a poorly characterized atypical DUSP widely conserved throughout evolution. Although no direct substrate has been firmly established, DUSP12 that has been implicated in protecting cells from stress, regulating ribosomal biogenesis, and modulating cellular DNA content. In this study, we used affinity- and proximity-based biochemical purification approaches coupled to mass spectrometry to identify the zinc finger protein ZPR9 as a novel DUSP12 interactor, which was validated by in-cell and *in-vitro* IP assays. Interestingly, ZPR9 binds to the unique zinc-binding domain of DUSP12, which previous reports indicated was important for many of DUSP12’s functions within the cell. Prior studies had implicated ZPR9 as a modulator of apoptosis, but it remained unclear if and how ZPR9 participated in the cell cycle and, more so, how it promoted cell death. Using mass spectrometry analyses, we found that overexpression of DUSP12 promoted de-phosphorylation of ZPR9 at Ser^143^. Overexpression of ZPR9, but not Ser^143^ phosphomimetic and phosphorylation-deficient mutants, led to an increase in pre-metaphase mitotic defects while knockdown of DUSP12 also showed mitotic defects in metaphase. Furthermore, knockdown of DUSP12 promoted, while knockdown of ZPR9 suppressed, stress-induced apoptosis. Our results support a model where DUSP12 protects cells from stress-induced apoptosis by promoting de-phosphorylation of ZPR9.

## INTRODUCTION

During tumorigenesis, quality control measures that supervise the fidelity of the cell cycle are often disrupted, leading to abnormal cell growth (1). In normal cells, cell cycle checkpoints are critical monitors of proper cell cycle progression and are, in turn, regulated by a complex network of signaling pathways (1). Among these mechanisms, protein phosphorylation serves an important role as a molecular switch to regulate events in signaling pathways that determine a cell’s fate, including cell cycle progression, cell division, and cell death (2, 3). While the regulatory role of kinases is well-established, less appreciated is the contribution of phosphatases in governing cell cycle progression (4).

Dual-specificity phosphatases (DUSPs), characterized by their unique ability to de-phosphorylate both tyrosine and threonine/serine residues within one substrate, are critical players of cell growth, survival, and death (5, 6). DUSPs de-phosphorylate and down-regulate the activity of mitogen-activated protein kinases (MAPKs) and phosphatidyl inositol 3-kinases (PI3Ks), which implies a role in regulating tumorigenesis through these signaling pathways (5, 6). Consistently, the dysregulation of DUSP activity is oncogenic or tumor-suppressive depending on the type of tumor (5). For example, DUSP1 was found to be overexpressed in pancreatic cancer and its downregulation decreased tumor formation in a pancreatic cancer mouse model (7). Similarly, the upregulation or overexpression of DUSP6 drives glioblastoma tumor formation in a glioblastoma mouse model (8). Therefore, as illustrated by the growing list of proliferative malignancies associated with deregulation of DUSP expression, DUSP phosphatases represent exciting new targets for study (5, 6).

DUSP12 is a member of the atypical DUSPs, named for their vastly diverse substrate specificity and function, and is highly conserved across mammalian species (5, 9). Several studies have linked aberrant expression of DUSP12 to the development of a wide range of cancers, highlighting the importance of investigating the biological activities of DUSP12 as dissecting its functions could provide insight into novel mechanisms supporting uncontrolled cell growth (10–14). Within atypical DUSPs, DUSP12 is unique in that it contains a C-terminal cysteine-rich zinc-binding domain (ZBD) in addition to an N-terminal phosphatase domain (15). Previous reports suggested that the ZBD mediates many of the observed effects of DUSP12 in the cell, including contributions to ribosome biogenesis, cell cycle progression, and cell survival (16–18). However, the physiological role of DUSP12 remains unclear.

In this study, we reveal that DUSP12 interacts with zinc finger protein 9 (ZPR9), which mediates apoptotic cell death through direct interaction with and activation by MPK38 and ASK1 in a phosphorylation-dependent manner (19–23). Due to the role of ZPR9 in cell death, it is interesting and necessary to determine the relationship between DUSP12 and cell death. We demonstrate that DUSP12 directly binds to ZPR9, promotes its dephosphorylation, and protects cells from stress-induced cell death.

## RESULTS

### DUSP12 is important for cell division and cell cycle progression

To evaluate the functional role of DUSP12 in cell division, we depleted HeLa cells of endogenous DUSP12 by RNAi (Fig. 1A) and performed a multiparametric analysis to monitor cell, chromosome, and mitotic spindle morphology during cell division by immunofluorescence (IF) microscopy (Fig. 1B). Knockdown of DUSP12 led to an increased percentage of cells with defects in chromosome alignment during metaphase (siControl = 12.4 ± 6.5 and siDUSP12 = 27.4 ± 2.5, *p* < 0.01) (Fig. 1C, D). To examine the role of DUSP12 on mitotic duration, we coupled DUSP12 depletion in HeLa-eBFP cells to live-cell time-lapse microscopy (Fig. 1E). This analysis showed that depletion of DUSP12 led to a significant increase in the time from chromosome condensation to chromosome segregation (siDUSP12 = 83.07 ± 54.46 min, p = 0.0004 compared with siControl = 58.47 ± 22.31 min) (Fig. 1F-H).

**Fig. 1.**
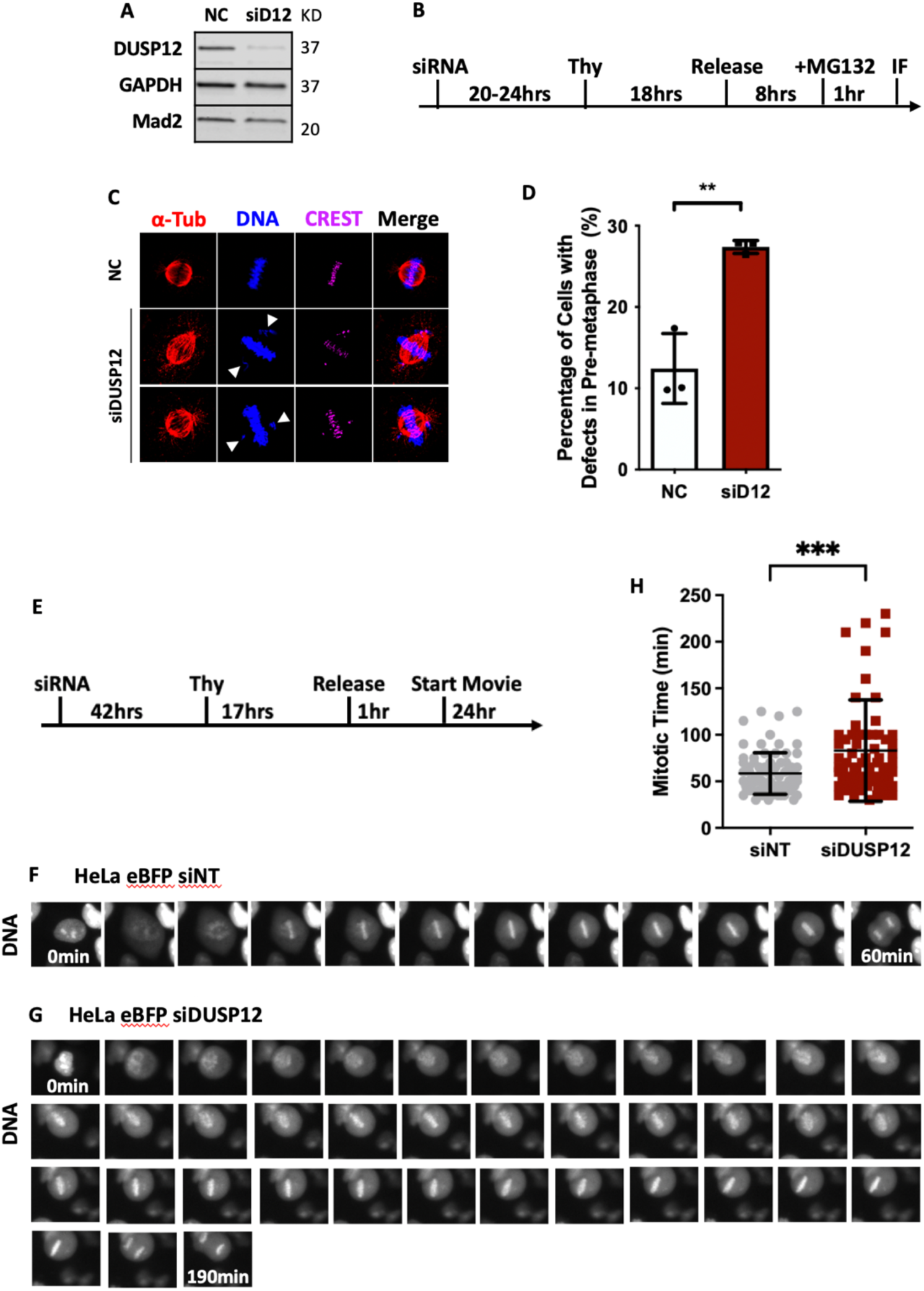
Knockdown of DUSP12 leads to metaphase defects. **A.** siRNA knockdown of endogenous DUSP12 in HeLa cells. **B.** Schematic of experiments performed in (C). **C.** Knockdown of DUSP12 leads to chromosome misalignment in metaphase. **D.** Quantification of cells with misaligned chromosomes shown in (C). **E.** Schematic of live-cell time-lapse microscopy experiment performed in (F) and (G). **F-G.** Live-cell time-lapse microscopy of HeLa-eBFP cells treated with negative control siRNA (F) or siDUSP12 (G) undergoing cell division. **H.** Quantification of the timing of mitosis from chromosome condensation to chromosome segregation (y-axis) for the indicated treatments.

In addition to the mitotic defects observed upon depleting DUSP12, a significant difference was observed in its cell cycle profile (Fig. S1A-E). Using flow cytometry, cells depleted of DUSP12 for 72 h accumulated in G2/M phase compared to control cells (Fig. S1E). To clarify how this affected cell cycle progression, we coupled HeLa fluorescence ubiquitination cell cycle indicator (FUCCI) cells, which exhibit color changes according to the phase of the cell cycle, to live-cell time-lapse IF microscopy. HeLa FUCCI cells were depleted of endogenous DUSP12 by RNAi (Fig. S1A). Cells were then synchronized with thymidine at G1/S phase and released into fresh media. Cells were imaged live four-hours post-release for 24 h. Both control and DUSP12-depleted conditions started at S/G2/M phase 4h post-thymidine release. However, as control cells continued to progress to G1 phase at 13 h post-thymidine release, DUSP12-depleted cells lagged in S/G2/M. This lagging in DUSP12-depleted cells persisted even 28 h post-thymidine release. Together, these results suggest that DUSP12 depletion leads to a slowing of cell division.

### DUSP12 directly interacts with and binds to ZPR9 via its unique zinc-binding domain

To further define DUSP12’s cellular role, we sought to determine its protein interactome in mitotic cells. We established inducible localization and affinity purification (LAP=EGFP-TEV-S-Peptide)-tagged and biotin identification 2 (BioID2)-tagged DUSP12 HeLa stable cell lines, which were used to express LAP-/BioID2-DUSP12. LAP affinity and BioID2 proximity biochemical purifications were then carried out and the eluates were analyzed by LC-MS/MS. The mass spectrometry data was analyzed and visualized as protein interaction/association networks with the CANVS software using the mitogen-activated protein kinase Gene Ontology (GO) terms (Supplementary Fig. S2A-C) (24). Both the interaction and association networks identified ZPR9 (aka ZNF622) as a novel DUSP12 interaction/association (Fig. 2A, B). To validate the DUSP12-ZPR9 interaction, we performed LAP-DUSP12 immunoprecipitations (IPs) and confirmed that ZPR9 co-IP’d with DUSP12 in both asynchronous cells (Fig. 2C) and Taxol-arrested mitotic cells (Fig. 2D) by immunoblot analysis.

**Fig. 2.**
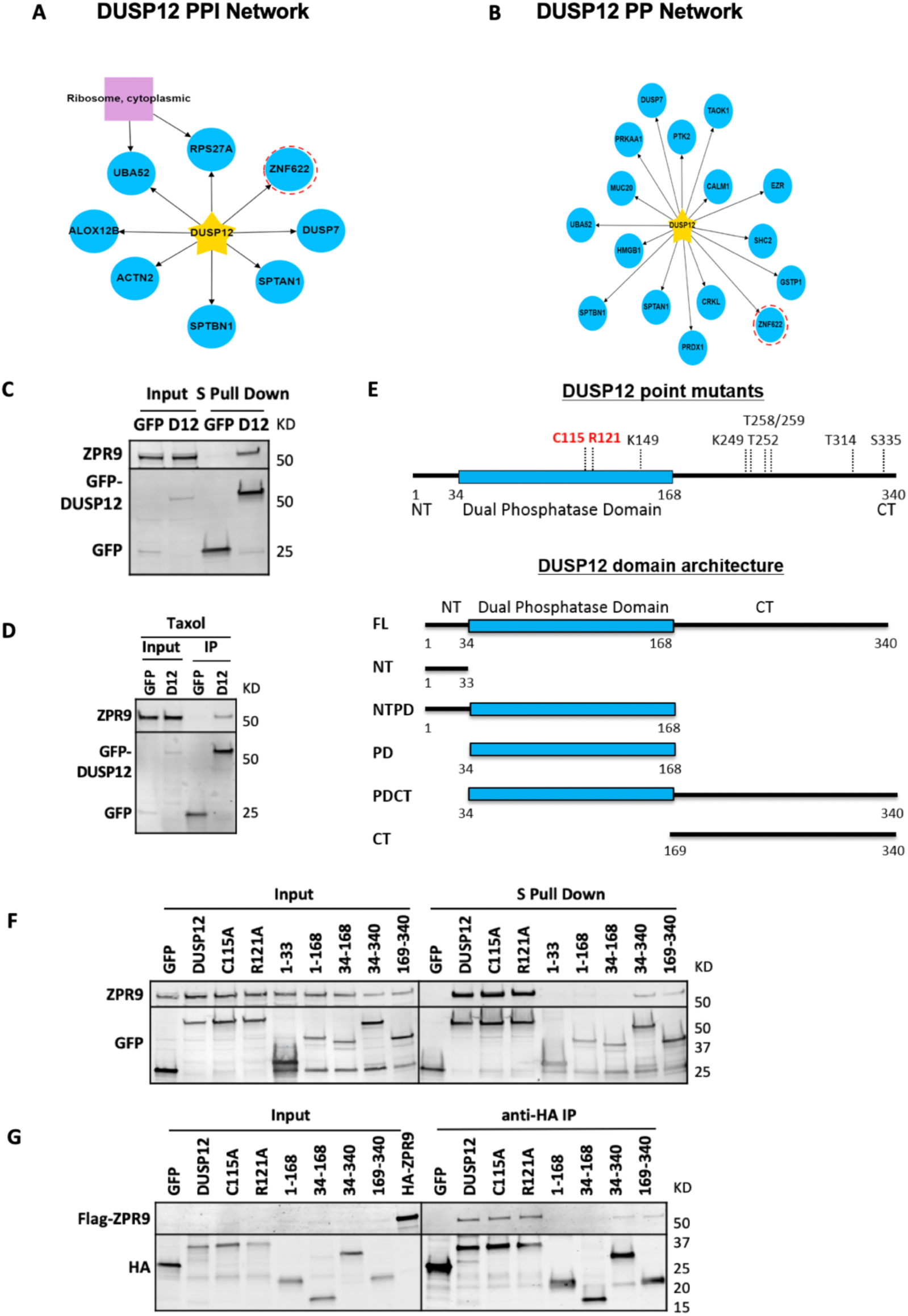
DUSP12 directly interacts with and binds to ZPR9 via its C-terminal Zinc Binding Domain. **A**, **B** Proteomic analysis of DUSP12 identifies ZPR9 as a novel interactor. **C**, **D** DUSP12 interacts with ZPR9 in asynchronized cells (**C**) and mitotic cells (**D**). **E** Schematic of DUSP12 point mutants and domain architecture. **F**, **G** ZPR9 binds to the C terminus of DUSP12 in HeLa cells (**F**) and *in vitro* (**G**), and the ZPR9-DUSP12 interaction is independent of DUSP12 phosphatase activity.

As the zinc-binding domain (ZBD) is important for many of DUSP12’s cellular functions, including its role in cell cycle regulation, we sought to determine whether the DUSP12 ZBD was important for binding to ZPR9. To do this in an unbiased manner, we generated a series of DUSP12 truncations (full length, N-terminus, N-terminus + phosphatase domain, phosphatase domain, phosphatase domain + C-terminus, and C-terminus) and catalytically dead mutants (C115A or R121A) (Fig. 2E and Fig. S3A-G). We then developed a series of LAP-DUSP12 inducible stable cell lines capable of expressing these DUSP12 truncations and mutations, and performed a series of IPs to investigate which domains of DUSP12 were involved in ZPR9 binding. These IPs showed that the N-terminus and the phosphatase domain of DUSP12 both failed to associate with ZPR9 (Fig. 2F). Whereas, DUSP12 full length, C-terminus, C-terminus + phosphatase, and the C115A and R121A mutants all efficiently pulled down ZPR9 (Fig. 2F). These results suggested that the DUSP12 N-terminus, phosphatase domain, and catalytic activity were dispensable for interacting with ZPR9, while the C-terminus was indispensable. To further verify this, we performed *in vitro* binding assays with FLAG-tagged ZPR9 and HA-tagged DUSP12 full-length, truncations, and catalytic dead mutants. Similar results were observed, where ZPR9 co-IP’d with DUSP12 full length, C-terminus, C-terminus + phosphatase, and the C115A and R121A mutants (Fig. 2G). Taken together, our results indicated that the DUSP12 C-terminus (amino acids 169-340), which contained the ZBD interacted directly with ZPR9.

### Overexpression of ZPR9 leads to mitotic defects

Since DUSP12 interacted with ZPR9 during mitosis (Fig. 2D), we evaluated whether ZPR9 was important for cell division and, further, how this related to its interaction with DUSP12. Immunoblot analyses showed that endogenous ZPR9 was expressed constitutively throughout the cell cycle, which was similar to the DUSP12 expression profile (Supplementary Fig. S4A-C). Consistent with the idea that DUSP12 and ZPR9 interact during cell division, IF microscopy showed that they shared similar localization patterns, particularly during mitosis (Supplementary Fig. S4D). More specifically, endogenous ZPR9 co-localized weakly to the spindles during prometaphase and metaphase, which was more apparent following overexpression of DUSP12 (Supplementary Fig. S4D, E).

To further assess the role of ZPR9 in cell division, we generated a LAP-ZPR9 stable cell line to analyze the consequences of ZPR9 overexpression on mitotic progression by IF microscopy (Fig. 3A-C). Overexpression of ZPR9 led to a pronounced increase in the percentage of mitotic cells with defects in metaphase (ZPR9 OE = 61.5 ± 2.5, p < 0.05 compared to NC = 30.3 ± 5.4) but not post-metaphase (Fig. 3D, F). These defective cells also led to a significant increase in mitotic cells with malformed spindles (multipolar and unfocused) (ZPR9 OE = 40 ± 3.5, p < 0.05 compared to NC = 10.3 ± 0.4) (Fig. 3C, E).

**Fig. 3.**
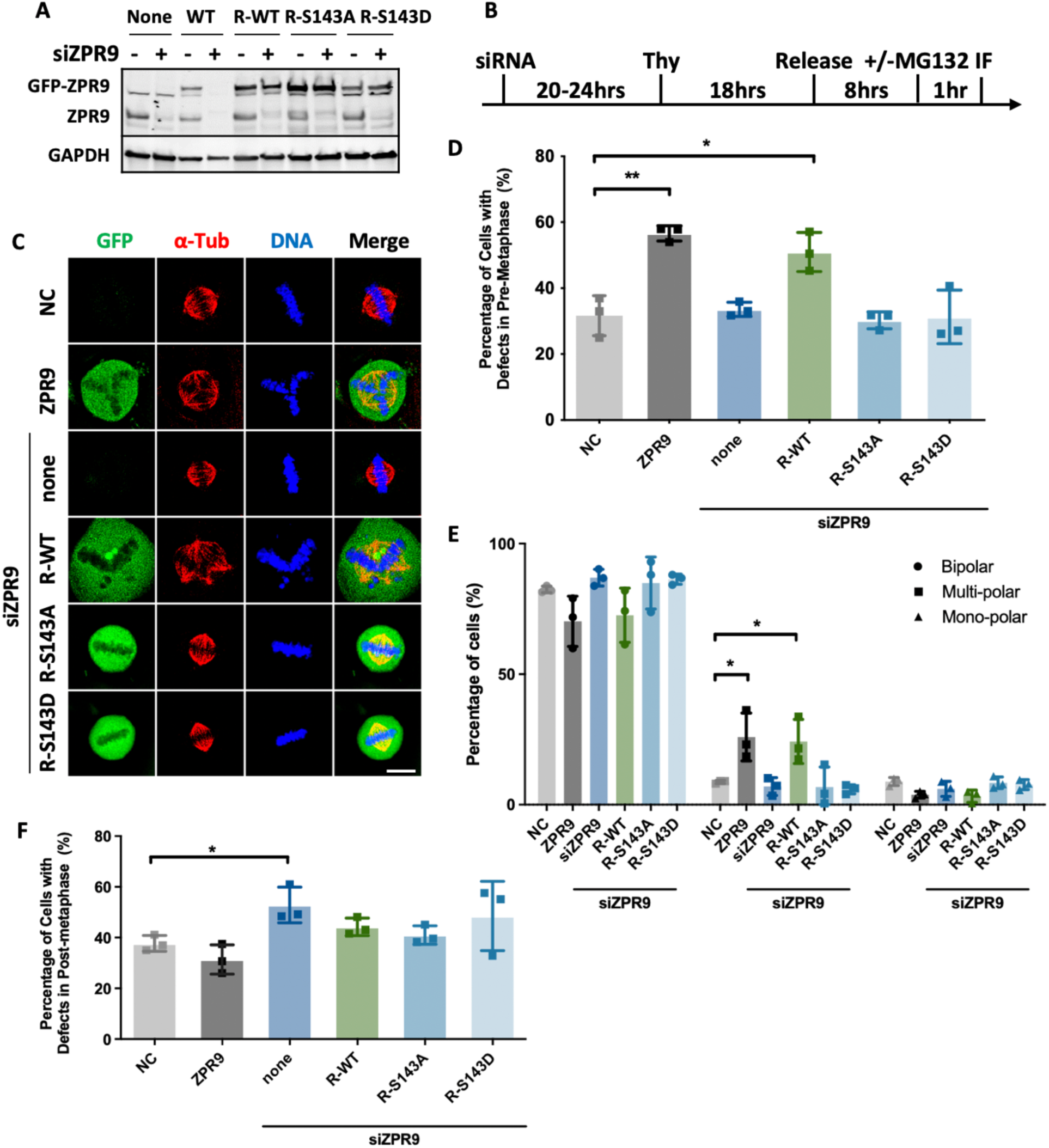
Overexpression of ZPR9 leads to mitotic defects. **A** siRNA knockdown of ZPR9 wild type but not siRNA resistant mutants. **B** Schematic of the immunofluorescence microscopy experiments performed in (C). **C** Overexpression of ZPR9 leads to metaphase defects in HeLa cells. Scale bars: 10 µm. **D**, **E** Quantification of cells with metaphase defects shown in **C**. **F** Quantification of cells with post-metaphase defects.

Given DUSP12’s capacity as a phosphatase, we next asked if DUSP12 modification of ZPR9 influenced cell division. From mass spectrometry analysis of inducible LAP-DUSP12 stable cells (Supplementary Fig. S5A-C), we identified S143 as a potential site of de-phosphorylation on ZPR9 by DUSP12. To evaluate the functionality of the identified phosphorylation site on ZPR9, phosphorylation-deficient (S/A) and phosphomimetic (S/E) mutations were introduced into LAP-ZPR9, and their effects were analyzed by IF. These mitotic defects were rescued by both siRNA resistant pGLAP1-ZPR9-S143 phosphosite mutants (Supplementary Fig. S3F, G) but not the siRNA resistant pGLAP1-ZPR9 (Supplementary Fig. S3E) expressed at near endogenous levels. Together, these results suggest that the ZPR9 S143 phosphorylation site modified by DUSP12 rescued chromosomal alignment and segregation defects.

### DUSP12 protects cells from apoptosis by regulating ZPR9 activity

As both DUSP12 and ZPR9 had been separately linked to cell death, we sought to investigate their relationship as it relates to cell death in response to stress agents. HeLa cells that had been depleted of DUSP12 or ZPR9 by RNAi or induced to overexpress DUSP12 or ZPR9 by doxycycline addition were subjected to the cytotoxic agents taxol or hydrogen peroxide (Fig. 4A, B). We then monitored cell viability with the CellTiter-Glo assay and apoptosis with Caspase-Glo 3/7 assay. Overexpression of DUSP12 and knockdown of ZPR9 effectively protected HeLa cells from taxol and hydrogen peroxide induced cell death (Fig. 4C-F, 5A-B). However, depletion of DUSP12 and overexpression of ZPR9 had the opposite effect of suppressing stress-induced cell death (Fig. 4C-F, 5A-B). These results were also observed in SW527 cells (Supplementary Fig. S6A-D). Taking this a step further, we observed a similar differential apoptotic response in HeLa cells upon coupling overexpression of DUSP12 or ZPR9 with a panel of cytotoxic agents, including staurosporine, etoposide, bortezomib, and colchine (Fig. 5C-D). To confirm that the marked decrease in cell viability of the oxidative stress-treated DUSP12 and ZPR9 transfectants was indeed due to apoptosis, we performed an Annexin V staining assay and analyzed the cells by flow cytometry. Similar results were obtained where overexpression of DUSP12 and knockdown of ZPR9 protected cells from cytotoxic agent induced cell death (Fig. 4G and Supplementary Fig. S7). Together, these results indicated that ZPR9 promoted and DUSP12 inhibited stress-induced apoptosis.

**Fig. 4.**
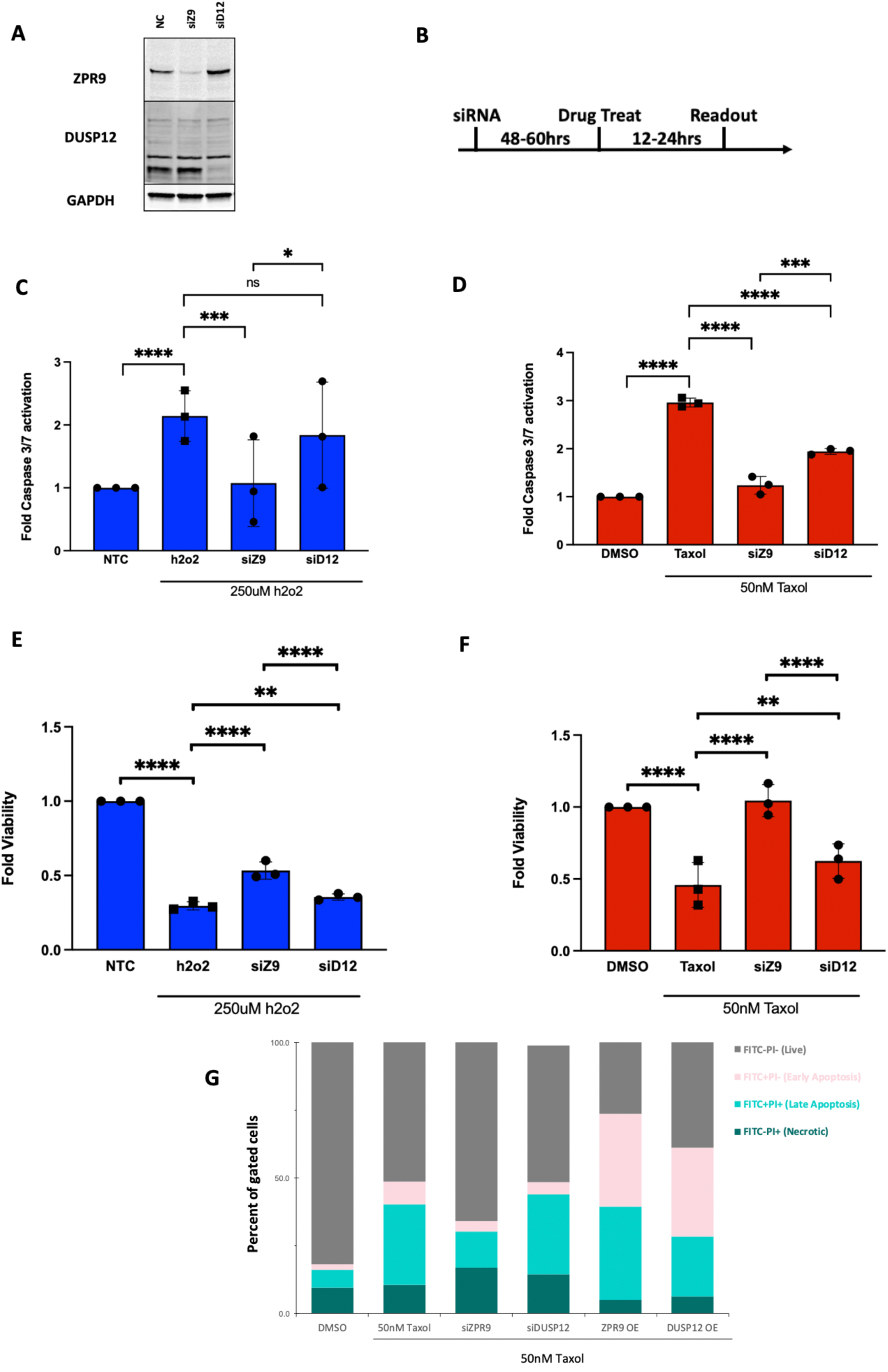
DUSP12 protects cells from apoptosis by regulating ZPR9. **A** siRNA knockdown or overexpression of DUSP12 and ZPR9 following 72 h and 36 h, respectively, in HeLa cells. **B** Schematic of experiments performed in **C-G**. **C-F** Genetically perturbed cells were coupled to cytotoxic agents (i.e. taxol or h2o2) after which viability was assessed with the CellTiter-Glo assay (**C**, **D**) and apoptosis with the Caspase-Glo assay (**E**, **F**), respectively, at 24 h for taxol and at 12 h for hydrogen peroxide. (*n*=3 technical replicates representative of at least *n*=3 independent experiments). **G** Apoptosis was assessed by staining treated cells with Propidium Iodide (PI) to distinguish live and dead cells and Annexin V to distinguish non- and pre-apoptotic cells (*n*=3 flow cytometric analyses of at least 10,000 cells, representative of *n*=3 independent experiments).

**Figure 5.**
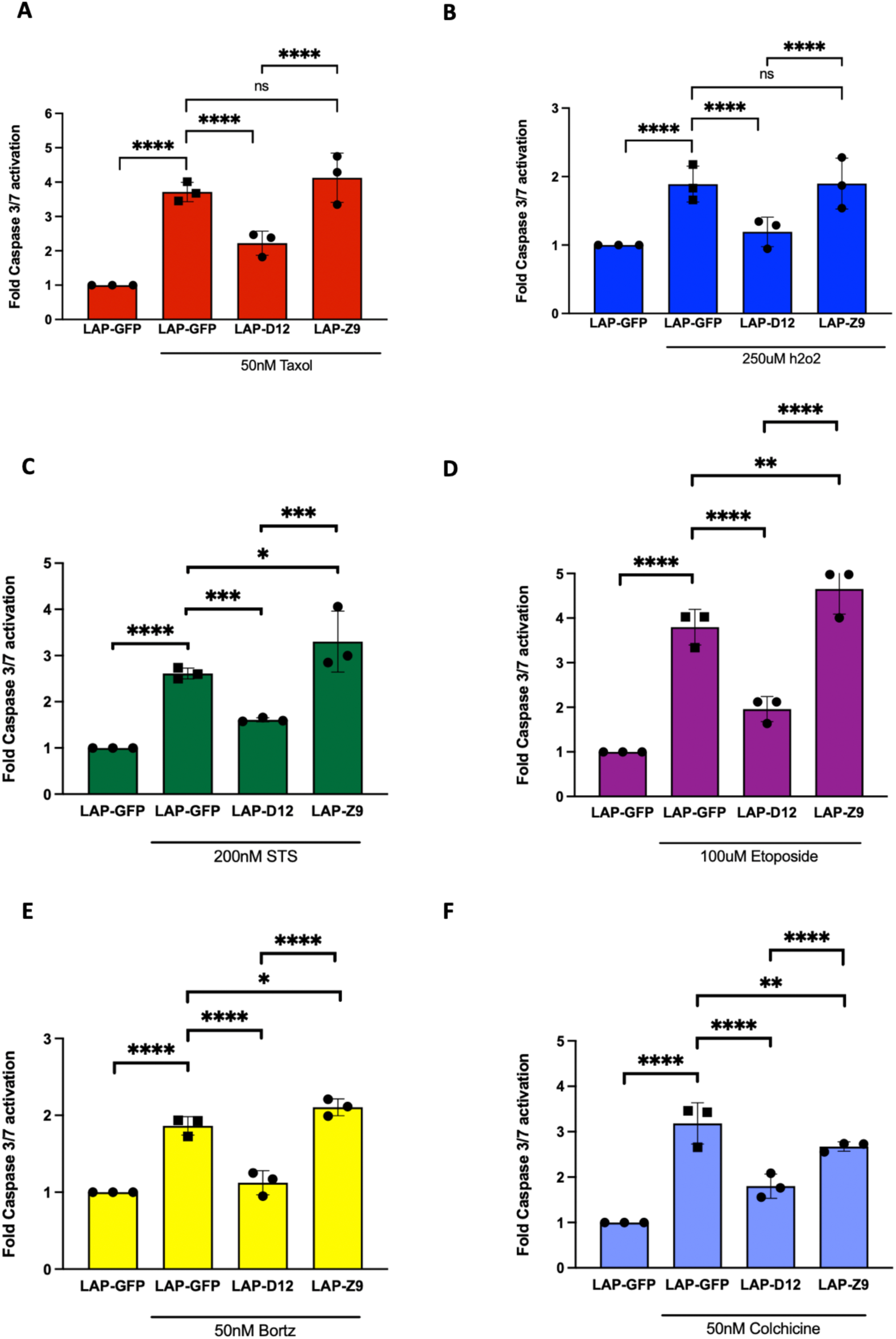
Overexpression of DUSP12 suppresses, while overexpression of ZPR9 promotes, stress-induced apoptosis. LAP-GFP, LAP-DUSP12, and LAP-ZPR9 HeLa stable cell lines were subjected to treatment with **A**) 50nM Taxol for 24 h, **B**) 250uM H_2_O_2_ for 12 h, **C**) 200nM Staurosporine for 24 h, **D**) 100uM Etoposide for 24h, **E**) 50nM Bortezomib for 24 h, and **F**) 50nM Colchicine for 24 h.

Next, we analyzed whether DUSP12 could suppress ZPR9 mediated cell death with or without the presence of cytotoxic agents. To test this, HeLa cells were transiently transfected with DUSP12 variants alone or with ZPR9 for 24h to evaluate proxy basal levels of cell death (Fig. 6A). While introduction of the full-length DUSP12 presented with levels of cell death lower than the control, expression of full-length ZPR9 displayed a clear increase in apoptotic cell death (Fig. 6A). Meanwhile, co-expression of ZPR9 and the zinc binding domain of DUSP12 displayed a marked reduction in apoptotic cell death that was comparable to DUSP12 alone (Fig. 6A). To further investigate whether DUSP12 can also protect cells exposed to stress conditions in which ZPR9 has been shown to play a cytotoxic role, LAP-GFP, LAP-DUSP12, and LAP-ZPR9 HeLa stable cell lines were subjected to transfection with an empty vector, full-length DUSP12, or ZPR9 and 50nM taxol for 24h. While expression of full-length ZPR9 promoted a significant increase in taxol mediated cell death, co-expression with DUSP12 rescued cells from either stressor (Fig. 6C). Together, these results suggest that DUSP12 mitigates ZPR9 mediated cell death.

**Figure 6.**
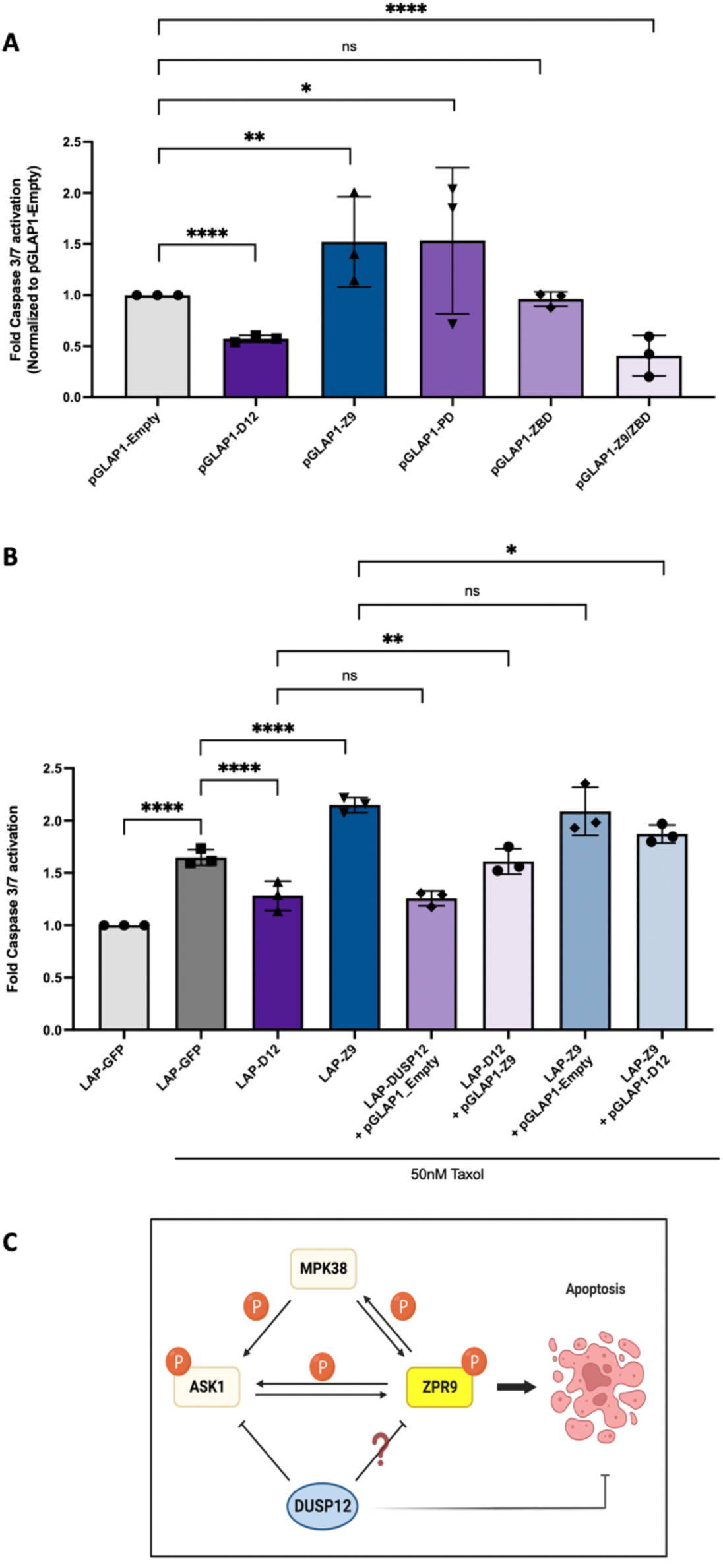
DUSP12 protects cells from apoptosis by regulating ZPR9. **A**. Caspase 3/7 levels were measured in HeLa cells transiently transfected with indicated constructs for 24h and left untreated. **B.** Caspase 3/7 levels were measured in LAP-GFP, LAP-DUSP12, and LAP-ZPR9 HeLa stable cell lines subjected to transfection with the indicated constructs and 50nM taxol for 24h. **C.** Potential model of how DUSP12 regulates ZPR9 activity to suppress cell death.

## DISCUSSION

This study advances our understanding of the factors and pathways that regulate cell cycle and cell death decisions. Our results are consistent with a model where DUSP12 binds to and counteracts the ability of ZPR9 to induce apoptosis in response to cytotoxic agents (Fig. 6C). We show that ZPR9 is a novel DUSP12 interacting partner and that this interaction relies on the DUSP12 zinc-binding domain. Further, DUSP12 promoted de-phosphorylation of ZPR9 at S143 and this modification was necessary to suppress ZPR9 mediated mitotic defects. Consistently, DUSP12 depletion led to mitotic defects in pre-metaphase and stress-induced cell death. Furthermore, overexpression of ZPR9 also led to a stress-induced cell death. Interestingly, co-expression of ZPR9/DUSP12 mitigated ZPR9-induced apoptosis. Together, these results establish the molecular crosstalk between DUSP12 and ZPR9 in the regulation of cell cycle progression and cell death.

Our results also help explain previous cancer studies where DUSP12 was found to be amplified or up-regulated in several types of cancers including hepatocellular carcinoma, neuroblastomas, sarcomas, and retinoblastomas, which in many cases correlated with disease progression (10, 13, 14, 25, 26). We show that DUSP12 overexpression protects cells from cell death in the cytotoxic effects of the chemotherapeutic agent taxol. Therefore, DUSP12 is a promising target in cancers that overexpress DUSP12 or harbor DUSP12 amplification. The development of specific DUSP12 pharmacological inhibitors will be critical to determining if inhibition of DUSP12 activity can render these cancers susceptible to cytotoxic agents through the activation of apoptosis, and whether combination treatments prove to be more efficacious than single treatments. Therefore, it will be of interest to determine if DUSP12 overexpression protects against other common types of chemotherapeutic agents that induce apoptosis or if DUSP12 has roles in protecting cells from other forms of cell death like autophagy or necrosis (27).

On the other hand, protection against apoptosis is important in certain stress scenarios, such as the protection of tissues and organs during trauma-driven apoptosis (28). Interestingly, recent studies have indicated that DUSP12 can protect against apoptosis in a hepatic ischemia-reperfusion injury mouse model (29, 30). Therefore, the development of pharmacological activators of DUSP12 and/or inhibitors of ZPR9 could prove to be beneficial in ameliorating liver failure during liver surgery. This reasoning could also be expanded to diseases that are driven by apoptosis, like neurodegenerative disorders, where the inhibition of apoptosis could ameliorate disease progression (28).

## DATA AVAILABILITY

The CANVS code used to analyze and visualize LC-MS/MS results was deposited at GitHub https://github.com/uclatorreslab/MassSpecAnalysis. The publicly available data for differential gene expression analysis and survival analysis were obtained from the online website GEPIA (http://gepia.cancer-pku.cn/index.html). The publicly available TCGA-BLCA cohort data (data includes 408 tumors and 19 normal samples) were obtained from the GDC Data Portal website (https://portal.gdc.cancer.gov/). All remaining data are contained within the article, Supplementary Information, or Original Data files.

## MATERIALS AND METHODS

### Cell culture

Table S1 lists all reagents and tools used in this study. HEK-293T, HeLa, HeLa Flp-In T-Rex, and SW527 cells were cultured in DMEM/Ham’s F-12 with L-glutamine (Genesee Scientific) supplemented with 10% fetal bovine serum and 1% penicillin-streptomycin (Life Technologies). Cells were incubated in 5% CO_2_ at 37°C and passaged twice weekly using 0.25% Trypsin-EDTA (Life Technologies). Modified HeLa Flp-In T-Rex and SW527 stable cell lines were generated as described below. Cells were tested for mycoplasma contamination using the MycoStrip mycoplasma detection kit (InvivoGen).

### Cell synchronization, transfection, and inhibitor treatment

For G1/S arrest and release experiments, cells were treated with 2mM thymidine (Sigma-Aldrich) for 18 h, washed three times with PBS, two times with complete media, and then released into fresh media. For G2/M arrests, cells were treated with 100nM taxol (Sigma-Aldrich) for 18 h. For metaphase arrests, cells were treated with 10uM MG132 (Millipore Sigma) for 1 h post-thymidine release. For siRNA experiments, cells were transfected with Silencer Select Validated siRNA (Thermo Fisher) against DUSP12, ZPR9, or a non-targeting control using RNAiMAX (Thermo Fisher) as described previously (31). For protein overexpression experiments, cells were transfected with 1ug LAP-tagged DUSP12 or ZPR9 using FuGENE HD (Promega) for the indicated times according to the manufacturer’s instructions. For combination experiments with cytotoxic agents (i.e. taxol or h2o2), cells were transfected and subsequently incubated with drugs for the indicated times.

### Plasmids, mutagenesis, and generation of stable cell lines

Site-directed mutagenesis using the QuikChange Lightning Site-Directed Mutagenesis Kit (Agilent) was performed to generate DUSP12 and ZPR9 mutants. cDNAs of GFP, DUSP12, DUSP12 truncations, DUSP12 catalytically dead mutants, ZPR9, and ZPR9 phospho-site mutants were cloned into pGLAP1, pGBioID2, pCS2-HA, or pCS2-FLAG via Gateway LR Clonase (Thermo Fisher) reactions. pGLAP1-only/DUSP12/DUSP12-C115A/DUSP12-R121A/DUSP12-truncations/ZPR9/ZPR9-S143A/ZPR9-S143D and pGBioID2-only/DUSP12 were used to generate doxycycline-inducible HEK-293T or HeLa Flp-In T-Rex LAP-GFP/DUSP12/DUSP12-C115A/DUSP12-R121A/DUSP12-truncations/ZPR9/ZPR9-S143A/ZPR9-S143D and BioID2-only/DUSP12 stable cell lines as described previously (32, 33).

### LAP/BioID2 purifications and LC-MS/MS analyses

For LAP affinity purifications, Taxol arrested LAP-tagged inducible stable cell lines were purified as previously described (32, 33). Briefly, LAP-only and LAP-DUSP12 stable cell lines were induced with 0.1 µg/mL doxycycline (Sigma-Aldrich) and arrested in mitosis with 100nM Taxol for 18 h before being harvested and lysed. Cleared lysates were subjected to tandem affinity purification by incubation with anti-GFP antibody beads, and the bound eluates were incubated with S-protein Agarose (Millipore Sigma). Final eluates were trypsinized for subsequent LC-MS/MS analysis. For BioID2 proximity purifications, biotinylated proteins were purified from Taxol arrested BioID2-tagged inducible stable cell lines as previously described (31, 34). Briefly, BioID2-only and BioID2-DUSP12 stable cell lines were pre-incubated with DMEM/Ham’s F-12 supplemented with 10% streptavidin Dynabead-treated (Thermo Fisher) FBS overnight at 4°C. Cells were induced with 0.1 µg/mL Dox and treated with 100 nM Taxol and 50 µM Biotin for 16 h before being harvested and lysed. Cleared lysates were purified by incubation with Dynabeads overnight at 4°C, and final eluates were trypsinized for downstream LC-MS/MS analysis. For mapping the dephosphorylation site on ZPR9 by DUSP12, LAP-only, LAP-ASK1, and LAP-DUSP12 were expressed with 0.1 µg/mL doxycycline for 18 h. The induced stable cell lines were harvested and lysed in LAP200 lysis buffer (50 mM Hepes pH 7.4, 200 mM KCl, 1 mM EGTA, 1 mM MgCl2, 10% glycerol) supplemented with 0.05% NP-40, 0.5 mM DTT and protease inhibitor cocktail (Thermo Fisher). Cell lysates were incubated with an anti-ZPR9 antibody (Santa Cruz Biotechnology) coupled to Protein A-coupled Sepharose beads (Bio-Rad) for 2 h before being prepared for LC-MS/MS analysis. Mass spectrometry analysis on all samples was performed on a Thermo Q Exactive Plus Orbitrap as described previously (31). Protein-protein interaction (affinity) and protein-protein association (proximity) data were analyzed and visualized with CANVS (24). CANVS integrated data from the Biological General Repository for Interaction Datasets (BioGRID v.3.5) (35) and the Comprehensive Resource of Mammalian Protein Complexes (CORUM v. 3.0) (36) to create networks that associated proteins based on cellular mechanisms by using Gene Ontology (GO) terms (37) and the RCytoscapeJS (38, 39) visualization package.

### Microscopy

Cell fixation and microscopy were carried out as described previously (31). Briefly, cells grown on glass coverslips in a 24-well plate were fixed with 4% paraformaldehyde and permeabilized with 0.2% Triton X-100/PBS. Subsequently, cells were blocked with IF buffer (1x PBS, 5% Fish Gelatin, 0.1% TritonX-100) before being stained with 0.5 µg/ml Hoechst 33342 and the indicated primary antibodies in IF buffer for 1 h at room temperature. After three PBS washes, cells were incubated with secondary antibodies in IF buffer for 30 min at room temperature and washed three times with PBS. Coverslips were mounted with ProLong Gold Antifade mounting solution (Invitrogen) on glass slides and sealed with nail polish. Representative cells were selected and imaged either on a Leica DMI6000 microscope (63x/1.40 NA oil objective, Leica AF6000 Analysis Package), a Leica MICA microscope (63x/1.40 NA oil objective, MICA Analysis Package), or an ImageXpress XL imaging system (Molecular Devices) and exported as TIFF files. Primary and secondary antibodies and their corresponding information can be found in Supplementary Table S1.

### IPs and binding assays

For immunoprecipitation, LAP-tagged induced stable cell lines or transiently transfected HeLa cells were harvested and lysed, then incubated with S-protein Agarose (Millipore Sigma) for 2 h at 4°C. After incubation, bound beads were washed three times with LAP100 buffer (50 mM Hepes pH 7.4, 100 mM KCl, 1 mM EGTA, 1 mM MgCl2, 10% glycerol) supplemented with 0.05% NP-40 and 0.5 mM DTT. For *in vitro* binding assays, HA-tagged GFP/DUSP12/DUSP12-C115A/DUSP12-R121A/DUSP12-truncations and FLAG-tagged ZPR9 were expressed in a Quick Coupled Transcription/Translation System (Promega) and incubated together with Anti-HA magnetic beads (MBL) for 1.5 h at 4°C. After incubation, bound beads were washed three times with LAP200 buffer (50 mM Hepes pH 7.4, 200 mM KCl, 1 mM EGTA, 1 mM MgCl2, 10% glycerol) supplemented with 0.05% NP-40 and 0.5 mM DTT. Final IP and IVT eluates were resolved by SDS-Page for immunoblot analysis.

### Immunoblotting

Cell protein extracts were prepared from cells grown in 6-well plates. After harvesting, cells were lysed in LAP200 lysis buffer (50 mM Hepes pH 7.4, 200 mM KCl, 1 mM EGTA, 1 mM MgCl2, 10% glycerol) supplemented with 0.3% NP-40, 1uM DTT, Phosphatase Inhibitor and Protease Inhibitor Cocktail tablets (Thermo Fisher). Cell extracts were cleared by spinning at 15000 RPM for 10 min at 4°C. Protein concentration was quantified by BCA analysis and 6x SDS reducing sample buffer was added. Processed total lysates were separated by SDS-PAGE and transferred to a PVDF membrane (EMD Millipore). Membranes were incubated in blocking buffer (PBS, 0.5% BSA, 0.05% Tween-20, 0.02% SDS, 0.05% Proclin) and then with the indicated primary and secondary antibodies to visualize protein levels. The imaging of immunoblots was performed with a LI-COR Odyssey Imaging system.

### Cell viability and apoptosis assays

Cell viability was measured at the indicated times following combination chemical treatment using the CellTiter-Glo Assay (Promega) and apoptosis was measured using the Caspase-Glo 3/7 Assay (Promega) as described previously (41). Experiments were performed in 96-well plates in triplicates that were averaged and normalized to the negative DMSO-matched control. For Annexin V apoptosis assays, cells were stained with Annexin V-FITC (Thermo Fisher) for 15 min at room temperature, washed with PBS, and incubated for 15 min at 4°C with Propidium Iodide (Thermo Fisher). Following a final wash, data acquisition was performed using the Attune NxT Flow Cytometer (Invitrogen). Data was analyzed using FlowJo (BD Biosciences), gating against smaller cellular debris and events consistent with more than one cell per droplet.

### Antibodies

Supplementary Table S1 lists all primary and secondary antibodies used in this study.

### Quantification and statistical analysis

For IF microscopy quantification, three independent experiments were performed with 100 cells counted per experiment (n=300). For live cell time-lapse microscopy quantification, three independent experiments were performed for each condition with 25 cells counted per experiment (n=75). All cell viability and apoptosis assays were performed as three biological replicates with three technical replicates each. The data was analyzed using unpaired Student’s t-test in Figure 1D, 1H, 3D-F, 4C-F, 5A-F, 6A, 6B, S1D, S1E, S6C, and S6D. P-values were calculated using an unpaired Student’s t-test between two groups. Data is judged to be statistically significant when P<0.05. Asterisks indicate statistical significance as * P<0.05, ** P<0.01, *** P<0.001. All data is presented as mean ± standard deviation. GraphPad Prism 10 was used for statistical analysis. Aivia AI Image Analysis Software was used to quantify cells at each cell cycle phase in Figure S1D. ModFit LT 6.0 was used to analyze the cell cycle profile in Figure S1E.

## AUTHOR CONTRIBUTIONS

M.A., X.G., I.R., E.F.V., W.C., A.A.G., J.P.W., and J.Z.T. performed experiments, discussed results, and wrote the paper.

### ACKNOWLEDGEMENTS

Fig. 6C was created using BioRender.com.

## FUNDING

Work was supported by NIH-NIGMS R35GM139539 (JZT); NIH-NIGMS T32GM145388 and NIH F31GM154466 (MA); a grant to the University of California, Los Angeles from the HHMI through the James H. Gilliam Fellowships for Advanced Study Program and an MBI Whitcome Fellowship (EFV); and NIH P30DK063491 (JPW). The content is solely the responsibility of the authors and does not necessarily represent the official views of the National Institutes of Health.

## AUTHOR CONTRIBUTIONS

M.A., X.G., I.R., E.F.V., W.C., A.A.G., J.P.W., and J.Z.T. performed experiments, discussed results and wrote the paper.

## COMPETING INTERESTS

The authors declare no competing interests.

## ADDITIONAL INFORMATION

## Abbreviations

DUSP: dual specificity phosphatase
LAP: localization and affinity purification
IP: immunoprecipitation
PI: phosphatase inhibitor
IF: immunofluorescence
OE: overexpression
WT: wild type
NC: negative control
D12: DUSP12
Thy: thymidine
STS: Staurosporine
Bortz: Bortezomib
NT: N terminus
CT: C terminus

## Notes

https://github.com/uclatorreslab/MassSpecAnalysis.

### Competing Interest Statement

The authors have declared no competing interest.

## REFERENCES

1. Matthews HK, Bertoli C, de Bruin RAM. Cell cycle control in cancer. Nat Rev Mol Cell Biol. 2022;23(1):74–88.

2. Ardito F, Giuliani M, Perrone D, Troiano G, Lo Muzio L. The crucial role of protein phosphorylation in cell signaling and its use as targeted therapy (Review). Int J Mol Med. 2017;40(2):271–80.

3. Ong JY, Bradley MC, Torres JZ. Phospho-regulation of mitotic spindle assembly. Cytoskeleton (Hoboken). 2020;77(12):558–78.

4. Novak B, Kapuy O, Domingo-Sananes MR, Tyson JJ. Regulated protein kinases and phosphatases in cell cycle decisions. Curr Opin Cell Biol. 2010;22(6):801–8.

5. Gao PP, Qi XW, Sun N, Sun YY, Zhang Y, Tan XN, et al. The emerging roles of dual-specificity phosphatases and their specific characteristics in human cancer. Biochim Biophys Acta Rev Cancer. 2021;1876(1):188562.

6. Zandi Z, Kashani B, Alishahi Z, Pourbagheri-Sigaroodi A, Esmaeili F, Ghaffari SH, et al. Dual-specificity phosphatases: therapeutic targets in cancer therapy resistance. J Cancer Res Clin Oncol. 2022;148(1):57–70.

7. Liao Q, Guo J, Kleeff J, Zimmermann A, Buchler MW, Korc M, et al. Down-regulation of the dual-specificity phosphatase MKP-1 suppresses tumorigenicity of pancreatic cancer cells. Gastroenterology. 2003;124(7):1830–45.

8. Messina S, Frati L, Leonetti C, Zuchegna C, Di Zazzo E, Calogero A, et al. Dual-specificity phosphatase DUSP6 has tumor-promoting properties in human glioblastomas. Oncogene. 2011;30(35):3813–20.

9. Rios P, Nunes-Xavier CE, Tabernero L, Kohn M, Pulido R. Dual-specificity phosphatases as molecular targets for inhibition in human disease. Antioxid Redox Signal. 2014;20(14):2251–73.

10. Ju G, Zhou T, Zhang R, Pan X, Xue B, Miao S. DUSP12 regulates the tumorigenesis and prognosis of hepatocellular carcinoma. PeerJ. 2021;9:e11929.

11. Orbach SM, DeVaull CY, Bealer EJ, Ross BC, Jeruss JS, Shea LD. An engineered niche delineates metastatic potential of breast cancer. Bioeng Transl Med. 2024;9(1):e10606.

12. Cain EL, Braun SE, Beeser A. Characterization of a human cell line stably over-expressing the candidate oncogene, dual specificity phosphatase 12. PLoS One. 2011;6(4):e18677.

13. Kresse SH, Berner JM, Meza-Zepeda LA, Gregory SG, Kuo WL, Gray JW, et al. Mapping and characterization of the amplicon near APOA2 in 1q23 in human sarcomas by FISH and array CGH. Mol Cancer. 2005;4:39.

14. Capasso M, Diskin SJ, Totaro F, Longo L, De Mariano M, Russo R, et al. Replication of GWAS-identified neuroblastoma risk loci strengthens the role of BARD1 and affirms the cumulative effect of genetic variations on disease susceptibility. Carcinogenesis. 2013;34(3):605–11.

15. Jeong DG, Wei CH, Ku B, Jeon TJ, Chien PN, Kim JK, et al. The family-wide structure and function of human dual-specificity protein phosphatases. Acta Crystallogr D Biol Crystallogr. 2014;70(Pt 2):421–35.

16. Muda M, Manning ER, Orth K, Dixon JE. Identification of the human YVH1 protein-tyrosine phosphatase orthologue reveals a novel zinc binding domain essential for in vivo function. J Biol Chem. 1999;274(34):23991–5.

17. Kozarova A, Hudson JW, Vacratsis PO. The dual-specificity phosphatase hYVH1 (DUSP12) is a novel modulator of cellular DNA content. Cell Cycle. 2011;10(10):1669–78.

18. Lo KY, Li Z, Wang F, Marcotte EM, Johnson AW. Ribosome stalk assembly requires the dual-specificity phosphatase Yvh1 for the exchange of Mrt4 with P0. J Cell Biol. 2009;186(6):849–62.

19. Seong HA, Manoharan R, Ha H. Coordinate Activation of Redox-Dependent ASK1/TGF-beta Signaling by a Multiprotein Complex (MPK38, ASK1, SMADs, ZPR9, and TRX) Improves Glucose and Lipid Metabolism in Mice. Antioxid Redox Signal. 2016;24(8):434–52.

20. Seong HA, Gil M, Kim KT, Kim SJ, Ha H. Phosphorylation of a novel zinc-finger-like protein, ZPR9, by murine protein serine/threonine kinase 38 (MPK38). Biochem J. 2002;361(Pt 3):597–604.

21. Seong HA, Jung H, Manoharan R, Ha H. Positive regulation of apoptosis signal-regulating kinase 1 signaling by ZPR9 protein, a zinc finger protein. J Biol Chem. 2011;286(36):31123–35.

22. Seong HA, Manoharan R, Ha H. Smad proteins differentially regulate obesity-induced glucose and lipid abnormalities and inflammation via class-specific control of AMPK-related kinase MPK38/MELK activity. Cell Death Dis. 2018;9(5):471.

23. Seong HA, Manoharan R, Ha H. Zinc finger protein ZPR9 functions as an activator of AMPK-related serine/threonine kinase MPK38/MELK involved in ASK1/TGF-beta/p53 signaling pathways. Sci Rep. 2017;7:42502.

24. Velasquez EF, Garcia YA, Ramirez I, Gholkar AA, Torres JZ. CANVS: an easy-to-use application for the analysis and visualization of mass spectrometry-based protein-protein interaction/association data. Mol Biol Cell. 2021:mbcE21050257.

25. Hirai M, Yoshida S, Kashiwagi H, Kawamura T, Ishikawa T, Kaneko M, et al. 1q23 gain is associated with progressive neuroblastoma resistant to aggressive treatment. Genes Chromosomes Cancer. 1999;25(3):261–9.

26. Gratias S, Schuler A, Hitpass LK, Stephan H, Rieder H, Schneider S, et al. Genomic gains on chromosome 1q in retinoblastoma: consequences on gene expression and association with clinical manifestation. Int J Cancer. 2005;116(4):555–63.

27. Green DR, Llambi F. Cell Death Signaling. Cold Spring Harb Perspect Biol. 2015;7(12).

28. Singh R, Letai A, Sarosiek K. Regulation of apoptosis in health and disease: the balancing act of BCL-2 family proteins. Nat Rev Mol Cell Biol. 2019;20(3):175–93.

29. Qiu T, Wang T, Zhou J, Chen Z, Zou J, Zhang L, et al. DUSP12 protects against hepatic ischemia-reperfusion injury dependent on ASK1-JNK/p38 pathway in vitro and in vivo. Clin Sci (Lond). 2020;134(17):2279–94.

30. Boldorini R, Clemente N, Alchera E, Carini R. DUSP12 acts as a novel endogenous protective signal against hepatic ischemia-reperfusion damage by inhibiting ASK1 pathway. Clin Sci (Lond). 2021;135(1):161–6.

31. Guo X, Ramirez I, Garcia YA, Velasquez EF, Gholkar AA, Cohn W, et al. DUSP7 regulates the activity of ERK2 to promote proper chromosome alignment during cell division. J Biol Chem. 2021;296:100676.

32. Torres JZ, Miller JJ, Jackson PK. High-throughput generation of tagged stable cell lines for proteomic analysis. Proteomics. 2009;9(10):2888–91.

33. Bradley M, Ramirez I, Cheung K, Gholkar AA, Torres JZ. Inducible LAP-tagged Stable Cell Lines for Investigating Protein Function, Spatiotemporal Localization and Protein Interaction Networks. J Vis Exp. 2016;118(118):54870.

34. Garcia YA, Velasquez EF, Gao LW, Gholkar AA, Clutario KM, Cheung K, et al. Mapping Proximity Associations of Core Spindle Assembly Checkpoint Proteins. J Proteome Res. 2021;20(7):3414–27.

35. Stark C, Breitkreutz BJ, Reguly T, Boucher L, Breitkreutz A, Tyers M. BioGRID: a general repository for interaction datasets. Nucleic Acids Res. 2006;34(Database issue):D535–9.

36. Giurgiu M, Reinhard J, Brauner B, Dunger-Kaltenbach I, Fobo G, Frishman G, et al. CORUM: the comprehensive resource of mammalian protein complexes-2019. Nucleic Acids Res. 2019;47(D1):D559–D63.

37. Ashburner M, Ball CA, Blake JA, Botstein D, Butler H, Cherry JM, et al. Gene ontology: tool for the unification of biology. The Gene Ontology Consortium. Nat Genet. 2000;25(1):25–9.

38. Shannon P, Markiel A, Ozier O, Baliga NS, Wang JT, Ramage D, et al. Cytoscape: a software environment for integrated models of biomolecular interaction networks. Genome Res. 2003;13(11):2498–504.

39. Franz M, Lopes CT, Huck G, Dong Y, Sumer O, Bader GD. Cytoscape.js: a graph theory library for visualisation and analysis. Bioinformatics. 2016;32(2):309–11.

40. Torres JZ, Ban KH, Jackson PK. A Specific Form of Phospho Protein Phosphatase 2 Regulates Anaphase-promoting Complex/Cyclosome Association with Spindle Poles. Mol Biol Cell. 2010;21(6):897–904.

41. Xia X, Lo YC, Gholkar AA, Senese S, Ong JY, Velasquez EF, et al. Leukemia Cell Cycle Chemical Profiling Identifies the G2-Phase Leukemia Specific Inhibitor Leusin-1. ACS Chem Biol. 2019;14(5):994–1001.

