## Supplemental Material for "DUSP12 promotes cell cycle progression and protects cells from cell death by regulating ZPR9"

**Table of contents:** **Page**

**SUPPLEMENTARY FIGURES** **S2 – S8**

**SUPPLEMENTARY TABLE S1** **S9-S14**

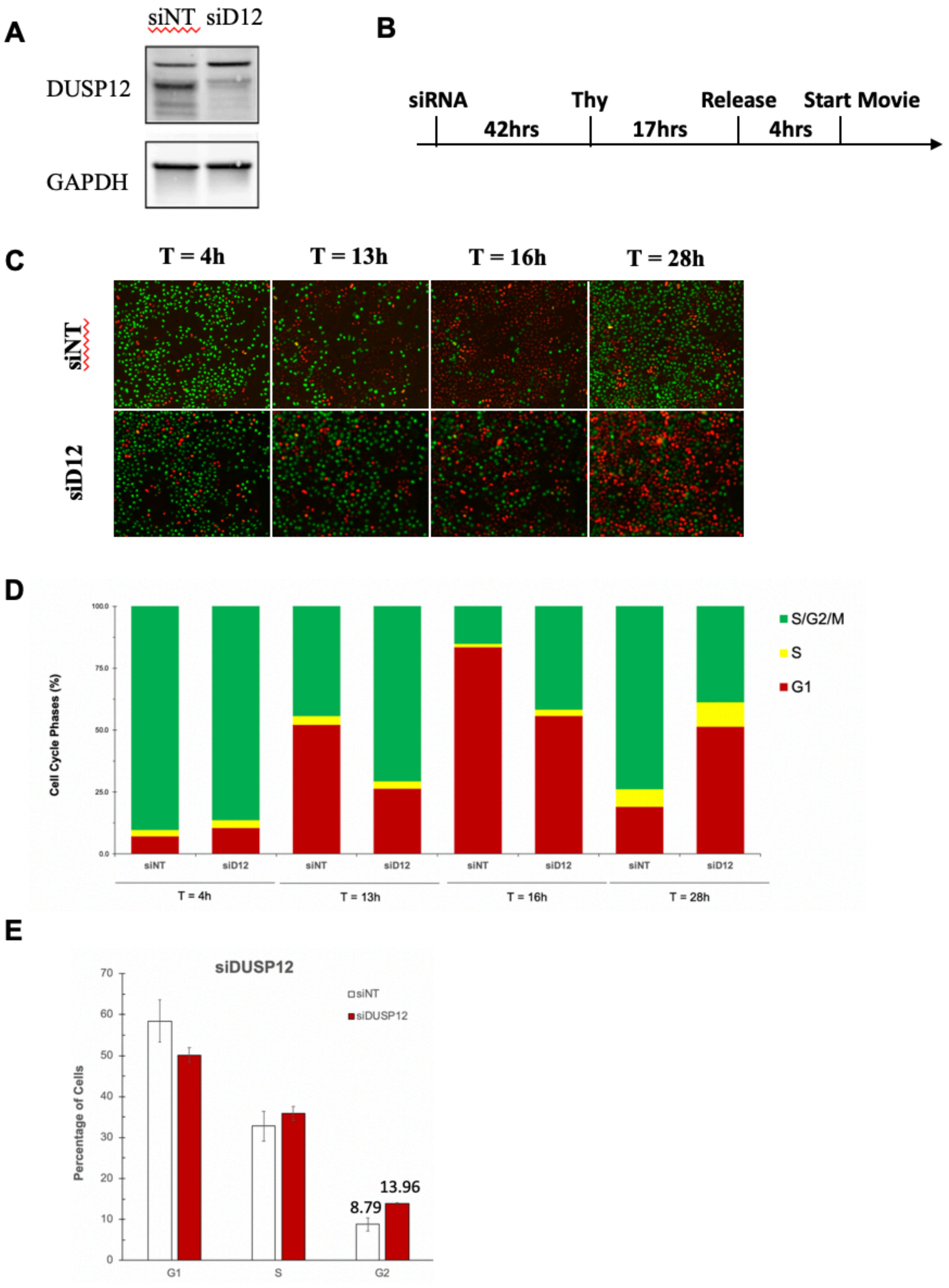
A
siNT siD12
DUSP12
GAPDH
B
siRNA
Thy
Release
Start Movie
42hrs
17hrs
4hrs
C
T = 4h
T = 13h
T = 16h
T = 28h
siNT
siD12
D
Cell Cycle Phases (%)
100.0
75.0
50.0
25.0
0.0
S/G2/M
S
G1
siNT
siD12
T = 4h
T = 13h
T = 16h
T = 28h
E
siDUSP12
Percentage of Cells
70
60
50
40
30
20
10
0
siNT
siDUSP12
8.79
13.96
G1
S
G2

**Supplementary Figure 1**. Knockdown of DUSP12 leads to mitotic slowing. **A.** siRNA knockdown of DUSP12 in HeLa FUCCI. **B.** Schematic of live-cell time-lapse microscopy experiment performed in (C). **C.** Knockdown of DUSP12 leads to S/G2/M arrest. Live-cell time-lapse microscopy of in HeLa FUCCI cells treated with negative control siRNA and siDUSP12 undergoing cell cycle phase changes **D.** Histograms show the percentage of cells in G 1 (red), early-S (yellow), or late-S/G 2 /M (green). Cells at each cell cycle phase were quantitatively assessed using Aivia AI Image Analysis Software. **E.** Knockdown of DUSP12 leads to G2/M arrest. Cell cycle profiling also shows that DUSP12 depletion arrested cells in G2/M phase.

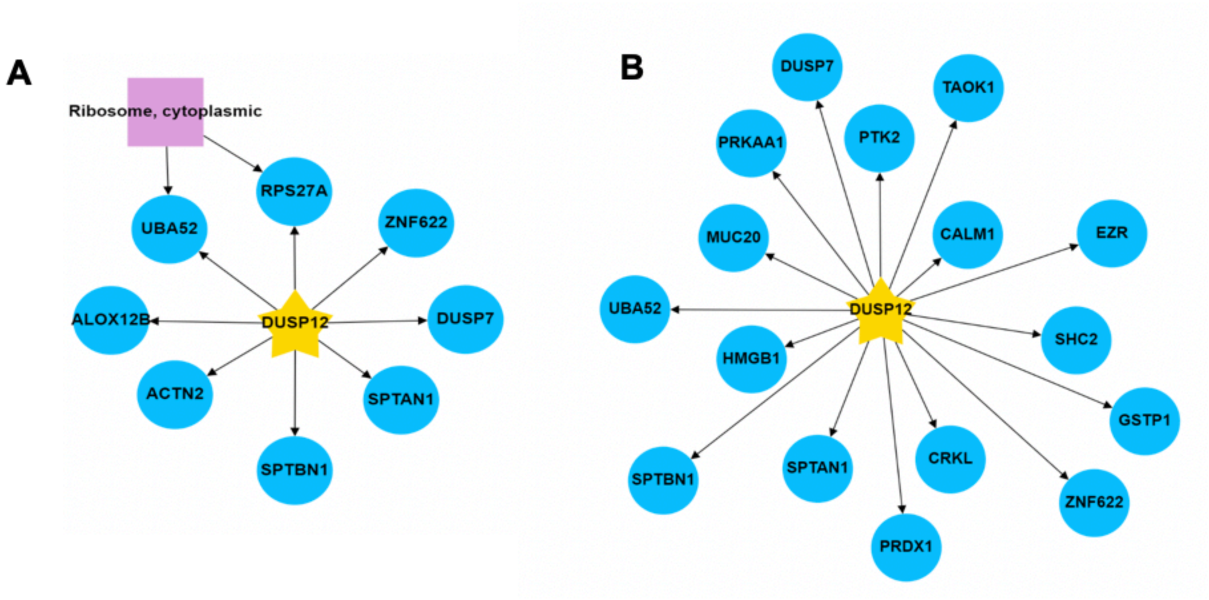


**C**

| GO:ID | Ontology |
|---|---|
| GO:0000165 | MAPK cascade |
| GO:0000186 | activation of MAPKK activity |
| GO:0000187 | activation of MAPK activity |
| GO:0000188 | inactivation of MAPK activity |
| GO:0032872 | regulation of stress-activated MAPK cascade |
| GO:0032873 | negative regulation of stress-activated MAPK cascade |
| GO:0043409 | negative regulation of MAPK cascade |
| GO:0043410 | positive regulation of MAPK cascade |
| GO:0051403 | stress-activated MAPK cascade |
| GO:1903753 | negative regulation of p38MAPK cascade |
| GO:0032874 | positive regulation of stress-activated MAPK cascade |

**Supplementary Figure S2. Proteomic analysis of the putative interactors of DUSP12**. **A** Mass spectrometry analysis of DUSP12 protein-protein interaction network. **B** Mass spectrometry analysis of DUSP12 proximity network. **C** Gene Ontology (GO) terms used to generate PPI and PP networks in Figure 1, **A** and **B**. Related to Figure 2.

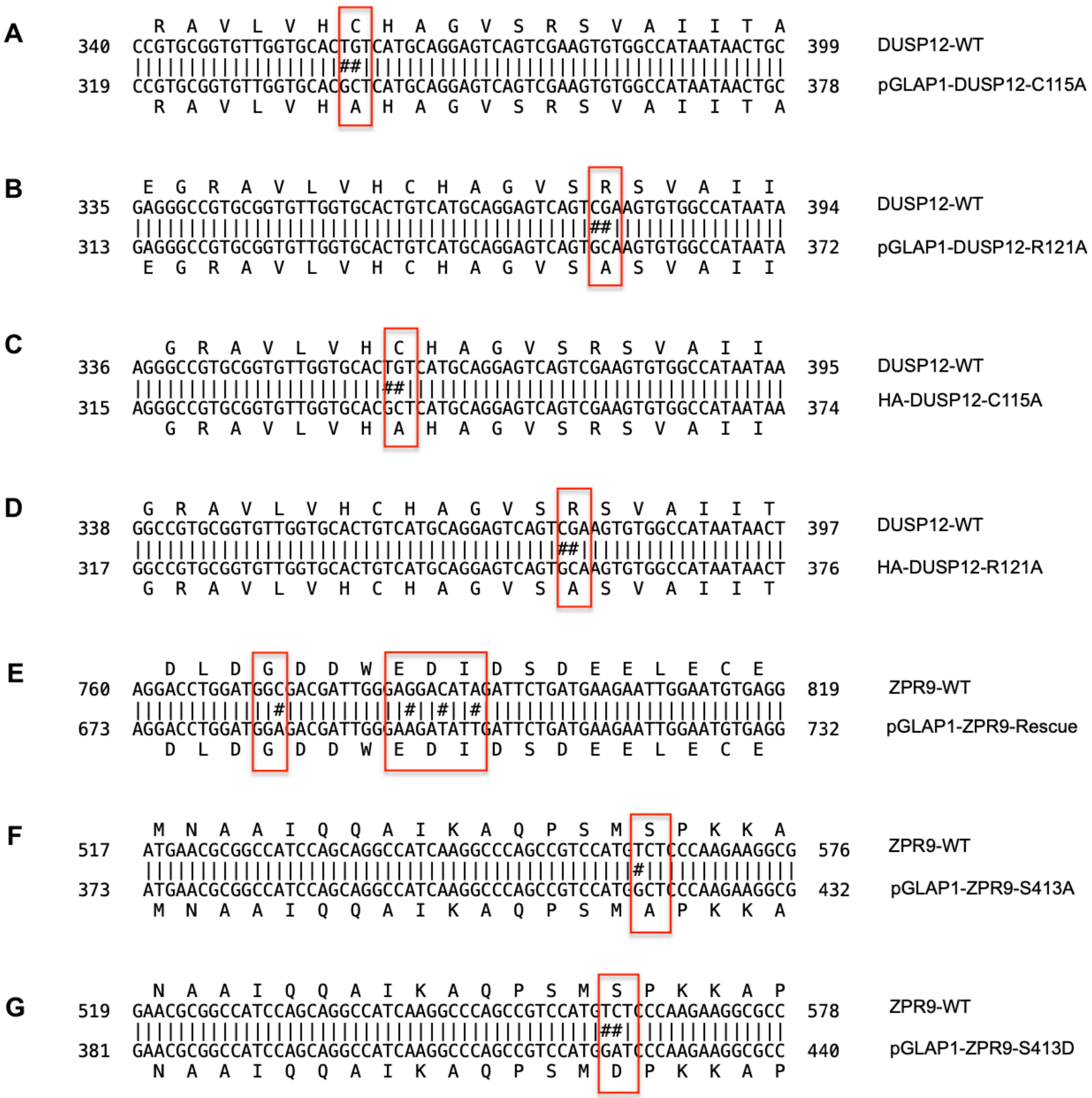


**Supplementary Figure S3. Verification of site-directed mutagenesis**. **A-G** Sequencing data of pGLAP1-DUSP12-C115A showing successful mutation of DUSP12-C115 into alanine (outlined by red box). **B** Sequencing data of pGLAP1-DUSP12-R121A showing successful mutation of DUSP12-R121 into alanine (outlined by red box). **C** Sequencing data of HA-DUSP12-C115A showing successful mutation of DUSP12-C115 into alanine (outlined by red box). **D** Sequencing data of HA-DUSP12-R121A showing successful mutation of DUSP12-R121 into alanine (outlined by red box). **E** Sequencing data of pGLAP1-ZPR9-Rescue showing successful mutation of ZPR9-WT into siRNA resistant mutant (highlighted in red boxes). **F** Sequencing data of pGLAP1-ZPR9-S143A showing successful mutation of ZPR9-S143 into alanine (outlined by red box). **G** Sequencing data of pGLAP1-ZPR9-S143D showing successful mutation of ZPR9-S143 into aspartate (outlined by red box). Related to Figure 2, Figure 3, and Figure 4.

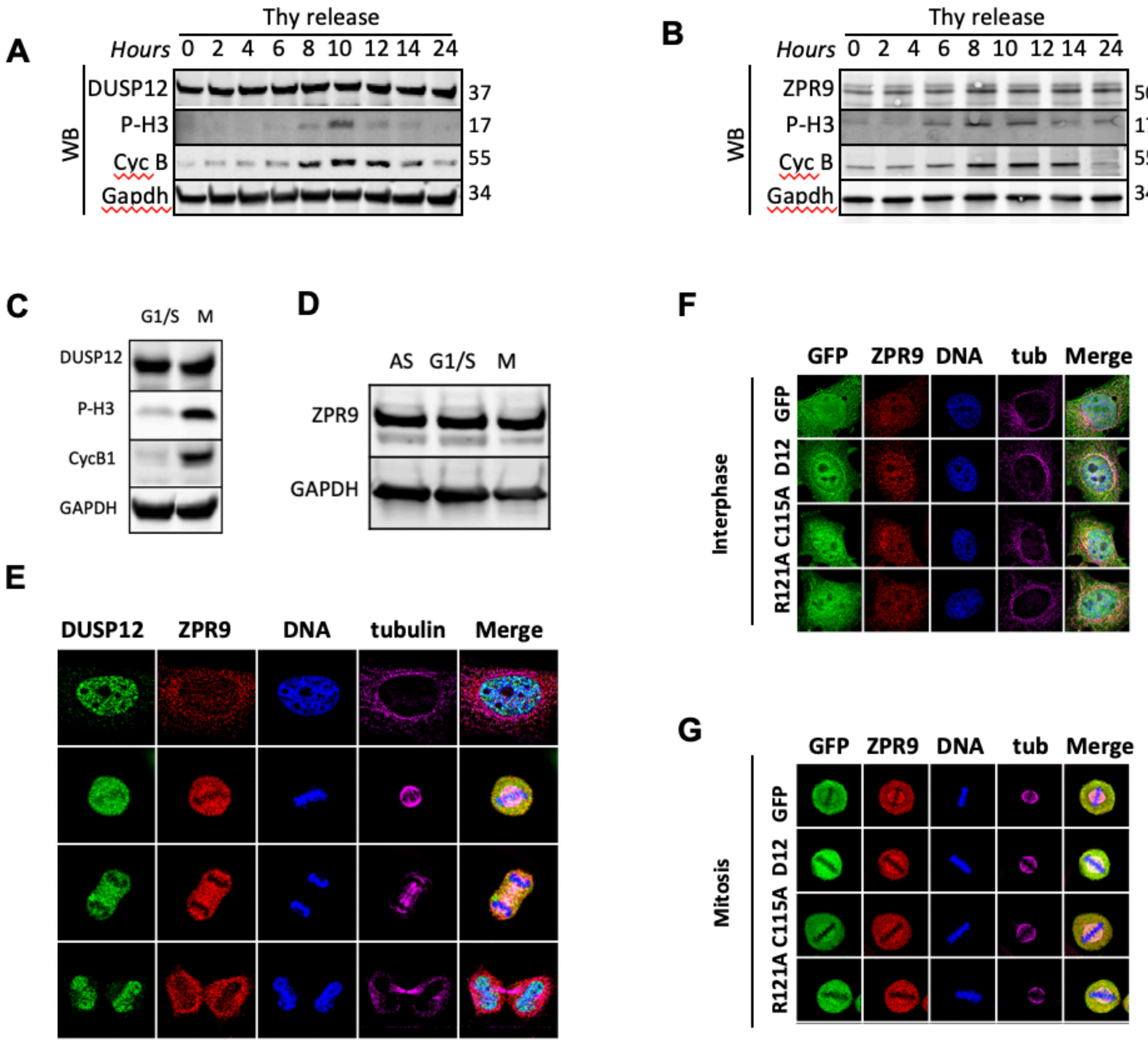


**Supplementary Figure 4.** Correlation of expression between DUSP12 and ZPR9. **A-D.** DUSP12 and ZPR9 are equally expressed throughout cell cycle. **E-G.** ZPR9 shares endogenous localization pattern of DUSP12 throughout the cell cycle. Related to Figure 3.

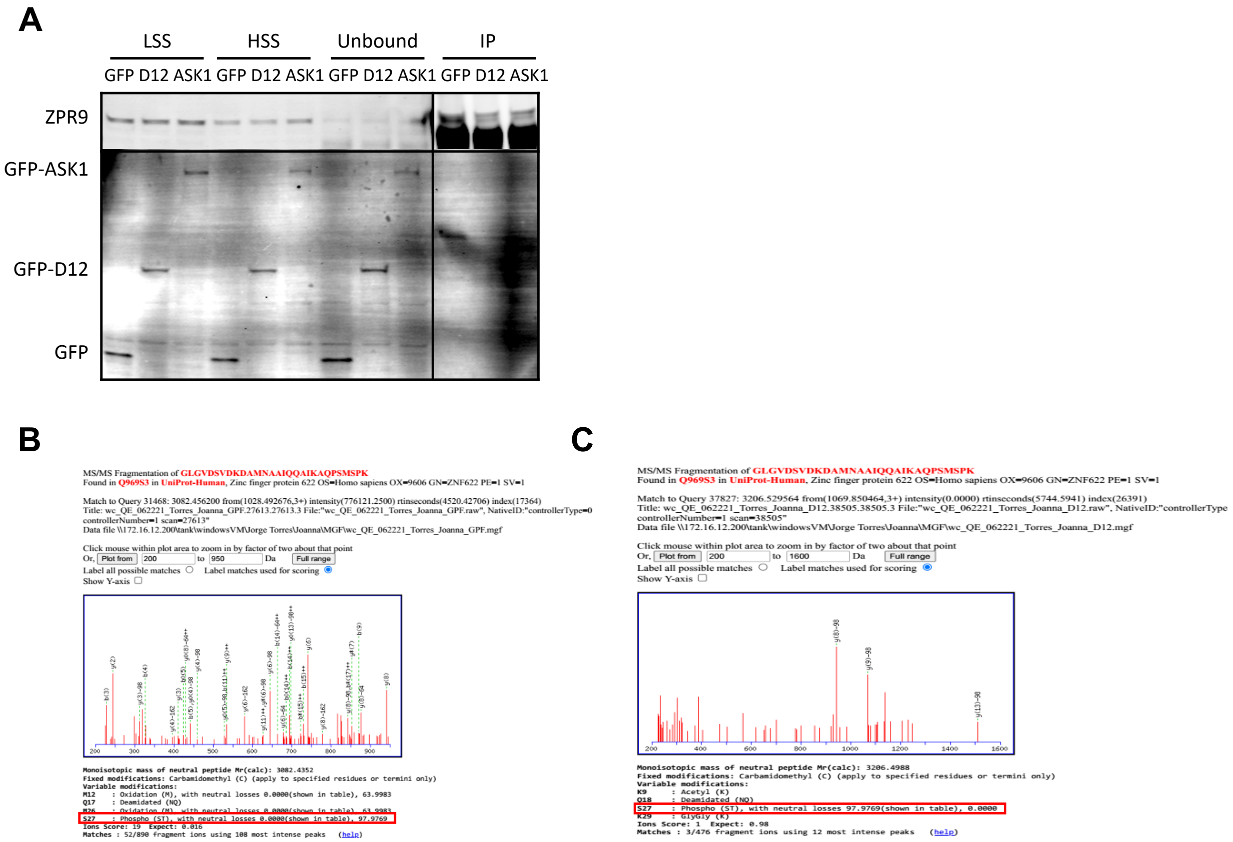


**Supplementary Figure S5**. **Identification of ZPR9 phosphorylation site using mass spectrometry**. **A** Anti-ZPR9 IP from HeLa LAP-only, LAP-DUSP12 and LAP-ASK1 cell lines for proteomics analysis shown in Figure 3. **B**, **C** Mass spectrometry spectra corresponding to LAP-only and LAP-DUSP12 peptide fingerprints generated with trypsin digestion. Related to Figure 3.

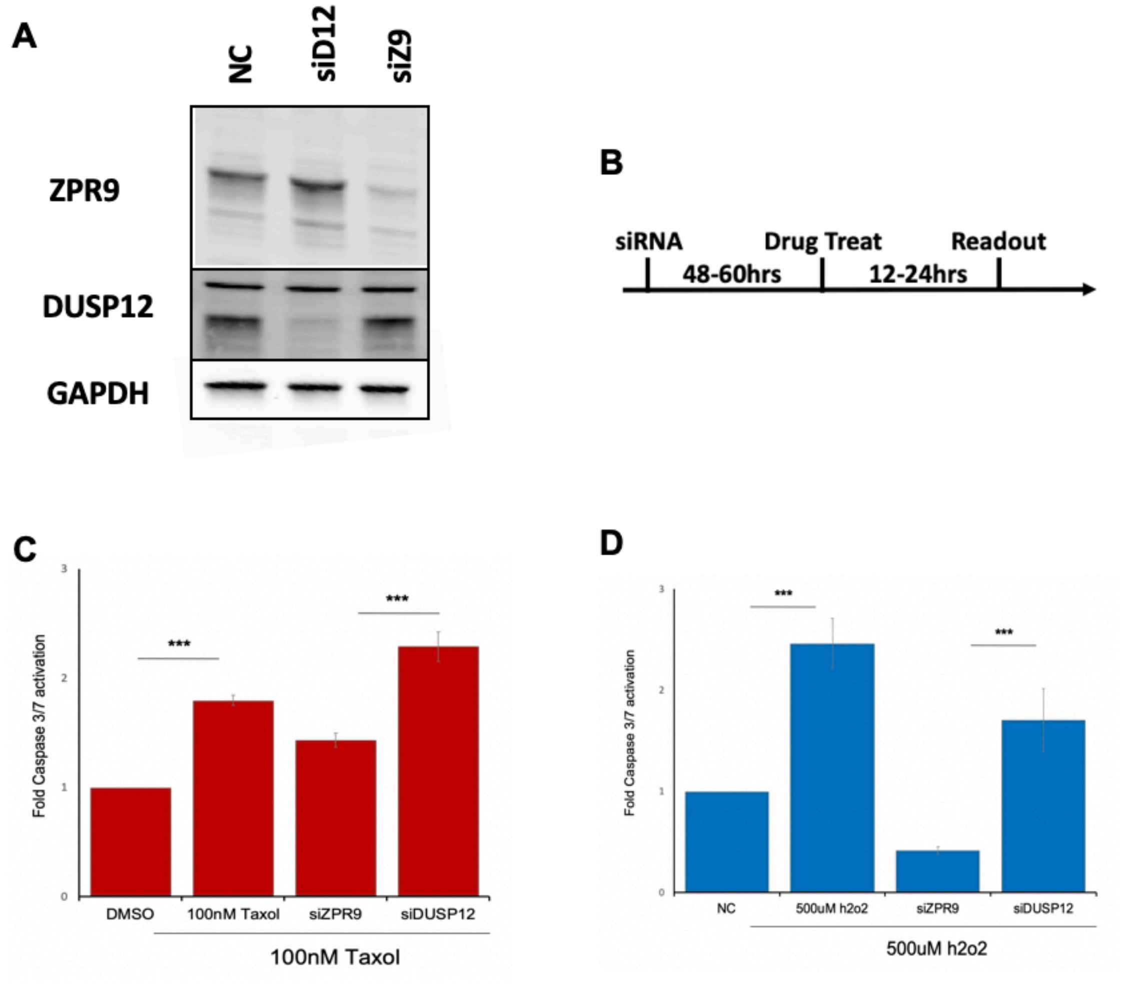


**Supplementary Figure S6. DUSP12 protects cells from apoptosis by regulating ZPR9**. **A** siRNA knockdown or overexpression of DUSP12 and ZPR9 following 72 h and 36 h, respectively, in SW527 cells. **B** Schematic of experiments performed in **C** and **D**. **C**, **D** Genetically perturbed cells were coupled to cytotoxic agents (i.e. taxol or h2o2) after which apoptosis was assessed by Caspase-Glo at 48 h for taxol and at 24 h for hydrogen peroxide. (N=3 technical replicates representative of at least N=3 independent experiments). Related to Figure 4.

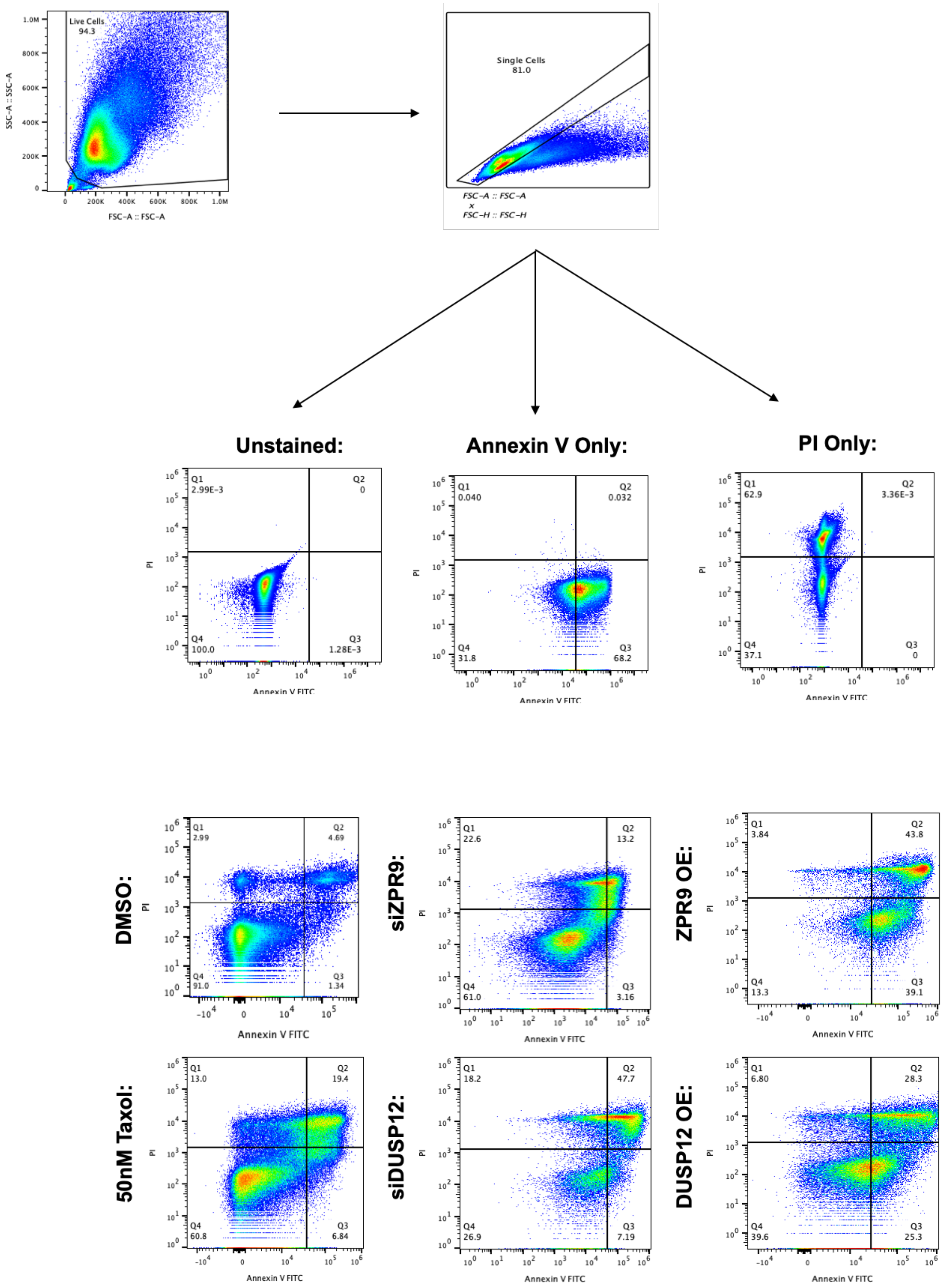


**Supplementary Figure S7**. **Example of the gating strategy used in the Annexin V experiments and representative flow plots**. Related to Figure 4.

**Table S1.** List of key reagents and resources used in this study.

| REAGENT or RESOURCE | SOURCE | IDENTIFIER |
|---|---|---|
| Antibodies | | |
| Rabbit polyclonal anti-DUSP12 | Proteintech | Cat# 15667-1-AP; RRID: AB_3085468 |
| Mouse monoclonal anti-ZNF622 | Santa Cruz Biotechnology | Cat# sc-100980; RRID: AB_2219571 |
| Rabbit polyclonal anti-ZPR9 Antibody | Bethyl Laboratories | Cat# A304-076A; RRID: AB_2621325 |
| Rabbit polyclonal anti-HA | Proteintech | Cat# 51064-2-AP; RRID: AB_11042321 |
| Mouse monoclonal anti-FLAG Dylight 800 Conjugated (clone 29E4.G7) | Rockland | Cat# 200-345-383; RRID: AB_10702994 |
| Human polyclonal anti-Centromere Protein/CREST | Antibodies Incorporated | Cat# 15-234-0001; RRID: AB_2687472 |
| Rat monoclonal anti-α-Tubulin (clone YOL1/34) | Bio-Rad | Cat# MCA78G; RRID: AB_325005 |
| Rabbit polyclonal anti-phospho-Histone H3 (Ser10) | Millipore | Cat# 06-570; RRID: AB_310177 |
| Goat polyclonal anti-Mad2 (C-19) | Santa Cruz Biotechnology | Cat# sc-6329; RRID: AB_648599 |
| Mouse monoclonal anti-GAPDH (clone 1E6D9) | Proteintech | Cat# 60004-1-Ig; RRID: AB_2107436 |
| Chicken polyclonal anti-GFP | Abcam | Cat# ab13970; RRID: AB_300798 |
| Donkey polyclonal anti-Rat IgG (H+L), Cy3 AffiniPure | Jackson ImmunoResearch | Cat# 712-165-153; RRID: AB_2340667 |
| Donkey polyclonal anti-Mouse IgG (H+L), Cy3 AffiniPure | Jackson ImmunoResearch | Cat# 715-165-151; RRID: AB_2315777 |
| Donkey polyclonal anti-Human IgG (H+L), Cy5 AffiniPure | Jackson ImmunoResearch | Cat# 709-175-149; RRID: AB_2340539 |
| Donkey polyclonal anti-Rat IgG (H+L), Cy5 AffiniPure | Jackson ImmunoResearch | Cat# 712-175-150; RRID: AB_2340671 |
| Donkey polyclonal anti-Chicken IgY (IgG) (H+L), Fluorescein (FITC) AffiniPure | Jackson ImmunoResearch | Cat# 703-095-155; RRID: AB_2340356 |
| Donkey polyclonal anti-Rabbit IgG (H+L), Fluorescein (FITC) AffiniPure | Jackson ImmunoResearch | Cat# 711-095-152; RRID: AB_2315776 |
| Donkey polyclonal anti-Goat IgG (H+L), IRDye 680RD | LI-COR Biosciences | Cat# 926-68074; RRID: AB_10956736 |
| Donkey polyclonal anti-Mouse IgG (H+L), IRDye 680RD | LI-COR Biosciences | Cat# 926-68072; RRID: AB_10953628 |
| Donkey polyclonal anti-Mouse IgG (H+L), IRDye 800CW | LI-COR Biosciences | Cat# 926-32212; RRID: AB_621847 |
| Donkey polyclonal anti-Chicken IgG (H+L), IRDye 800CW | LI-COR Biosciences | Cat# 926-32218; RRID: AB_1850023 |
| Donkey polyclonal anti-Rabbit IgG (H+L), IRDye 680RD | LI-COR Biosciences | Cat# 926-68073; RRID: AB_10954442 |

| | | |
|---|---|---|
| Donkey polyclonal anti-Rabbit IgG (H+L), IRDye 800CW | LI-COR Biosciences | Cat# 926-32213; RRID: AB_621848 |
| Chemicals, Peptides, and Recombinant Proteins | | |
| Paclitaxel | Sigma-Aldrich | Cat# T7191; CAS: 33069-62-4 |
| Hydrogen Peroxide | Sigma-Aldrich | Cat# 323381; CAS: 7722-84-1 |
| Staurosporine | Selleck Chemicals | Cat# S1421; CAS: 62996-74-1 |
| Etoposide | Selleck Chemicals | Cat# S1225; CAS: 33419-42-0 |
| Bortezomib | Selleck Chemicals | Cat# S1013; CAS: 179324-69-7 |
| Colchicine | Selleck Chemicals | Cat# S2284; CAS: 64-86-8 |
| Halt Protease Inhibitor Cocktail | Thermo Fisher Scientific | Cat# 87786 |
| Halt Phosphatase Inhibitor | Thermo Fisher Scientific | Cat# 78429 |
| Thymidine | Sigma-Aldrich | Cat# T1895; CAS: 50-89-5 |
| Doxycycline | Sigma-Aldrich | Cat# D9891; CAS: 24390-14-5 |
| MG132 | Millipore Sigma | Cat# 474790; CAS: 133407-82-6 |
| Hoechst 33342 | Thermo Fisher Scientific | Cat# H1399; CAS: 23491-52-3 |
| ProLong Gold Antifade Mountant | Thermo Fisher Scientific | Cat# P36934 |
| Lipofectamine RNAiMAX | Thermo Fisher Scientific | Cat# 13778150 |
| FuGENE HD Transfection Reagent | Promega | Cat# E2311 |
| FuGENE 6 Transfection Reagent | Promega | Cat# E2691 |
| S-protein Agarose | Millipore Sigma | Cat# 69704-4 |
| Anti-FLAG M2 magnetic beads | Sigma-Aldrich | Cat# M8823 |
| Biotin | Sigma-Aldrich | Cat# B4501-1G |
| Dynabeads MyOne Streptavidin C1 | Thermo Fisher Scientific | Cat# 65002 |
| Affi-Prep Protein A Resin | Bio-Rad | Cat# 156-0005 |
| Critical Commercial Assays | | |
| Gateway LR Clonase II Enzyme mix | Thermo Fisher Scientific | Cat# 11791020 |
| Gateway BP Clonase II Enzyme mix | Thermo Fisher Scientific | Cat# 11789020 |
| QuikChange Lightning Site-Directed Mutagenesis Kit | Agilent | Cat# 210518 |
| QIAprep Spin Miniprep Kit | QIAGEN | Cat# 27106 |
| PureYield Plasmid Midiprep System | Promega | Cat# A2495 |

| | | |
|---|---|---|
| SP6 TnT Quick Coupled Transcription/Translation System | Promega | Cat# L2080 |
| CellTiter-Glo 2.0 Cell Viability Assay | Promega | Cat# G9242 |
| Caspase-Glo 3/7 Assay System | Promega | Cat# G8091 |
| eBioscience Annexin V Apoptosis Detection Kit FITC | Thermo Fisher Scientific | Cat# 88-8005-74 |
| MycoStrip Mycoplasma Detection Kit | InvivoGen | Cat# REP-MYS-10 |
| Deposited Data | | |
| Affinity-based mass spectrometry performed with LAP-DUSP12 | This paper | |
| Proximity-based mass spectrometry performed with BioID2-DUSP12 | This paper | |
| Experimental Models: Cell Lines | | |
| SW527 cell line | ATCC | Cat# CRL-7940; RRID: CVCL_3799 |
| HeLa cell line | ATCC | Cat# CCL-2; RRID: CVCL_0030 |
| HeLa Flp-In T-Rex cell lines | Stephen Taylor Lab | (Tighe et al., 2004) |
| Inducible HeLa LAP-DUSP12 stable cell line | This paper | N/A |
| Inducible HeLa BioID2-DUSP12 stable cell line | This paper | N/A |
| Inducible HeLa LAP-DUSP12-C115A stable cell line | This paper | N/A |
| Inducible HeLa LAP-DUSP12-C115A stable cell line | This paper | N/A |
| Inducible HeLa LAP-DUSP12-R121A stable cell line | This paper | N/A |
| Inducible HeLa LAP-DUSP12-1-33aa stable cell line | This paper | N/A |
| Inducible HeLa LAP-DUSP12-1-168 stable cell line | This paper | N/A |
| Inducible HeLa LAP-DUSP12-34-168 stable cell line | This paper | N/A |
| Inducible HeLa LAP-DUSP12-34-340 stable cell line | This paper | N/A |
| Inducible HeLa LAP-DUSP12-169-340 stable cell line | This paper | N/A |
| Inducible HeLa LAP-ZPR9 stable cell line | This paper | N/A |
| Inducible HeLa LAP-ZPR9-rescue stable cell line | This paper | N/A |
| Inducible HeLa LAP-ZPR9-rescue-S143A stable cell line | This paper | N/A |
| Inducible HeLa LAP-ZPR9-rescue-S143D stable cell line | This paper | N/A |
| Oligonucleotides | | |
| siRNA against Non-Targeting | Thermo Fisher Scientific | Cat# 4390844 |

| | | |
|---|---|---|
| siRNA against DUSP12 | Thermo Fisher Scientific | Cat# 4390826; siRNA ID: s22244 |
| siRNA against ZNF622 | Thermo Fisher Scientific | Cat# 4392422; siRNA ID: s40388 |
| Primer for cloning DUSP12-C115A: Fwd 5'-TGACTCCTGCATGAGCGTGCACCAACACCG-3' | Eurofins Genomics | N/A |
| Primer for cloning DUSP12-C115A: Rev 5'-CGGTGTTGGTGCACGCTCATGCAGGAGTCA-3' | Eurofins Genomics | N/A |
| Primer for cloning DUSP12-R121A: Fwd 5'-TTATGGCCACACTTGCACTGACTCCTGCAT-3' | Eurofins Genomics | N/A |
| Primer for cloning DUSP12-R121A: Rev 5'-ATGCAGGAGTCAGTGCAAGTGTGGCCATAA-3' | Eurofins Genomics | N/A |
| Primer for cloning DUSP12-1-33aa: Fwd 5'-GGGGACAAGTTTGTACAAAAAAGCAGGCTTCATGGGGTTGGAGGCTCCGGGCCCG -3' | Eurofins Genomics | N/A |
| Primer for cloning DUSP12-1-33aa: Rev 5'-GGGGACCACTTTGTACAAGAAAGCTGGGTCTCATCCTGGCTGCACTTCCAGC-3' | Eurofins Genomics | N/A |
| Primer for cloning DUSP12-1-168aa: Fwd 5'-GGGGACAAGTTTGTACAAAAAAGCAGGCTTCATGGGGTTGGAGGCTCGGGCCCG -3' | Eurofins Genomics | N/A |
| Primer for cloning DUSP12-1-168aa: Rev 5'-GGGGACCACTTTGTACAAGAAAGCTGGGTCTCACATTGCCTGGTATAATTTCAGTTGC -3' | Eurofins Genomics | N/A |
| Primer for cloning DUSP12-34-168aa: Fwd 5'-GGGGACAAGTTTGTACAAAAAAGCAGGCTTCATGGGGTGTATTTCGGTGGGGCCGCG-3' | Eurofins Genomics | N/A |
| Primer for cloning DUSP12-1-34-168aa: Rev 5'-GGGGACCACTTTGTACAAGAAAGCTGGGTCTCACATTGCCTGGTATAATTTCAGTTGC -3' | Eurofins Genomics | N/A |
| Primer for cloning DUSP12-34-340aa: Fwd 5'-GGGGACAAGTTTGTACAAAAAAGCAGGCTTCATGGGGTGTATTTCGGTGGGGCCGCG-3' | Eurofins Genomics | N/A |
| Primer for cloning DUSP12-34-340aa: Rev 5'-GGGGACCACTTTGTACAAGAAAGCTGGGTCTCATATTTTTCCTGTTTGTGATCCCAAACAG-3' | Eurofins Genomics | N/A |
| Primer for cloning DUSP12-169-340aa: Fwd 5'-GGGGACAAGTTTGTACAAAAAAGCAGGCTTCATGGGGATACGAAGTGGATACCTCTAGTGCAA -3' | Eurofins Genomics | N/A |
| Primer for cloning DUSP12-169-340aa: Rev 5'-GGGGACCACTTTGTACAAGAAAGCTGGGTCTCATATTTTTCCTGTTTGTGATCCCAAACAG-3' | Eurofins Genomics | N/A |
| Primer for cloning ZPR9 S143A: Fwd 5'-CCAGCCGTCCATGGCTCCCAAGAAGGC-3' | Eurofins Genomics | N/A |
| Primer for cloning ZPR9 S143A: Rev 5'-GCCTTCTTGGGAGCCATGGACGGCTGG -3' | Eurofins Genomics | N/A |

| | | |
|---|---|---|
| Primer for cloning ZPR9 S143D: Fwd 5'-CCCAGCCGTCCATGGATCCCAAGAAGGCGC -3' | Eurofins Genomics | N/A |
| Primer for cloning ZPR9 S143D: Rev 5'-GCGCCTTCTTGGGATCCATGGACGGCTGGG -3' | Eurofins Genomics | N/A |
| Primer for cloning ZPR9-Rescue: Fwd 5'-GACCTGGATGGCGACGATTGGGAGGACATAGATTCTGAT-3' | Eurofins Genomics | N/A |
| Primer for cloning ZPR9-Rescue: Rev 5'-ATCAGAATCTATGTCCTCCCAATCGTCGCCATCCAGGTC-3' | Eurofins Genomics | N/A |
| Recombinant DNA | | |
| pDONR221-DUSP12 | DNASU Plasmid Repository | Clone ID: HsCD00041750 |
| pLX304-ZNF622 | DNASU Plasmid Repository | Clone ID: HsCD00441669 |
| pDONR221-MAP3K5 | DNASU Plasmid Repository | Clone ID: HsCD00860258 |
| pGLAP1-DUSP12 | This paper | N/A |
| pGLAP1-DUSP12-C115A | This paper | N/A |
| pGLAP1-DUSP12-R121A | This paper | N/A |
| pGLAP1-DUSP12-1-33aa | This paper | N/A |
| pGLAP1-DUSP12-1-168aa | This paper | N/A |
| pGLAP1-DUSP12-34-168aa | This paper | N/A |
| pGLAP1-DUSP12-34-340aa | This paper | N/A |
| pGLAP1-DUSP12-169-340aa | This paper | N/A |
| pGBioID2-DUSP12 | This paper | N/A |
| pCS2-HA-DUSP12 | This paper | N/A |
| pCS2-HA-DUSP12 | This paper | N/A |
| pCS2-HA-DUSP12-C115A | This paper | N/A |
| pCS2-HA-DUSP12-R121A | This paper | N/A |
| pCS2-HA-DUSP12-1-33aa | This paper | N/A |
| pCS2-HA-DUSP12-1-168aa | This paper | N/A |
| pCS2-HA-DUSP12-34-168aa | This paper | N/A |
| pCS2-HA-DUSP12-34-340aa | This paper | N/A |
| pCS2-HA-DUSP12-169-340aa | This paper | N/A |
| pGLAP1-ZPR9 | This paper | N/A |
| pGLAP1-ZPR9-S143A | This paper | N/A |
| pGLAP1-ZPR9-S143D | This paper | N/A |
| pGLAP1-ZPR9-rescue | This paper | N/A |
| pGLAP1-ZPR9-rescue-S143A | This paper | N/A |
| pGLAP1-ZPR9-rescue-S143D | This paper | N/A |
| pCS2-Flag-ZPR9 | This paper | N/A |
| pGLAP1-GFP | This paper | N/A |
| pCS2-HA-GFP | This paper | N/A |
| Software and Algorithms | | |
| Adobe Illustrator | Adobe | http://shop.adobe.com/store/adbehap/DisplayHomePage |

| | | |
|---|---|---|
| GraphPad Prism 5 | GraphPad | RRID: SCR_002798 |
| Biorender | Biorender | RRID: SCR_018361 |
